# Loss of PKN2 drives fibroblast reprogramming and extracellular matrix remodelling in pulmonary fibrosis

**DOI:** 10.64898/2026.09.02.748906

**Authors:** Catherine E McMullan, Oscar A. Peña, Poojitha Rajasekar, Anita Valand, Panayiota Stylianou, Gill Elliott, Molly Whitfield, Mathuscha Ratnasingham, Sahana Narayan, Alfie J Hall, Beatriz Goncalves, Vijay Mistry, Liesl Carr, Hilary Marshall, Bin Liu, Donald J L Jones, Peter Bradding, Richard J Allen, Louise V Wain, Rachel L Clifford, Colleen B Maxwell, Katy M Roach

## Abstract

**Introduction:** Idiopathic pulmonary fibrosis (IPF) is a progressive fibrotic lung disease characterised by aberrant fibroblast function, extracellular matrix (ECM) remodelling and defective tissue repair. Protein kinase N2 (PKN2) is associated with accelerated forced vital capacity decline in IPF, but its functional role in pulmonary fibrosis remains unknown. We hypothesised that PKN2 regulates fibroblast phenotype and tissue repair.

**Methods:** PKN2 expression was assessed in human lung tissue, induced sputum and primary airway and parenchymal fibroblasts from non-fibrotic controls and patients with interstitial lung disease (ILD). DNA methylation was profiled using the Illumina HumanMethylationEPIC array. PKN2 function was investigated by siRNA-mediated depletion in primary human lung fibroblasts using transcriptomic, proteomic and functional analyses. Tissue repair was assessed following pharmacological PKN inhibition in zebrafish.

**Results:** PKN2 expression was reduced in ILD lung tissue and primary airway and parenchymal fibroblasts and further suppressed by TGF-β1. Differential methylation was identified across the PKN2 locus in both fibroblast populations. Integrated transcriptomic and proteomic profiling following PKN2 depletion revealed coordinated remodelling of ECM, cell adhesion, non-canonical WNT and VEGF pathways, including dysregulation of COL1A1, WNT, VEGF and MMP1. PKN2 loss increased VEGF and MMP-1 secretion and accelerated fibroblast wound closure. PKN inhibition altered epithelial organisation and collagen fibre alignment during zebrafish wound repair.

**Conclusion:** PKN2 loss drives fibroblast reprogramming and aberrant ECM remodelling, establishing PKN2 as an important regulator of pulmonary fibroblast homeostasis and tissue repair.

## Introduction

Idiopathic pulmonary fibrosis (IPF) is a chronic, progressive and ultimately fatal interstitial lung disease (ILD) characterised by irreversible extracellular matrix (ECM) deposition and destruction of the normal alveolar architecture. IPF affects approximately three million people worldwide and carries a median survival of only 3–5 years following diagnosis^1,2^. Although antifibrotic therapies, including nintedanib, nerandomilast and pirfenidone, slow disease progression, they do not reverse established fibrosis^3–6^.

IPF is characterised by excessive scarring of the lung parenchyma and the accumulation of ECM components. The disease arises from repetitive microinjury to the alveolar epithelial cells which is followed by a dysregulated repair process, including increased activation of fibroblasts ^7^. Fibroblasts play a central role in the synthesis, secretion, and maintenance of ECM components, thereby supporting key physiological processes including tissue homeostasis and wound repair ^7,8^. In IPF, aberrant fibroblast activation is accompanied by excessive deposition of pro-fibrotic ECM proteins, notably collagen and fibronectin. The accumulation of fibrotic tissue progressively remodels the lung parenchyma, narrowing airway spaces, reducing lung compliance and capacity, increasing the diffusion distance for gas exchange, and disrupting normal alveolar architecture. Collectively, these structural changes impair respiratory function and are associated with marked declines in quality of life, mental health, and overall survival ^9–11^. As fibrosis advances, progressive loss of pulmonary function ultimately culminates in respiratory failure. Although the precise initiating cause of IPF remains unknown, epidemiological and genetic studies implicate several risk factors, including aging, smoking, occupational exposures and genetic susceptibility ^12–18^. Current evidence suggests that disease development most likely arises from a complex interplay between environmental exposures and genetic predisposition^15,19^

Despite recent advances in understanding fibrotic signalling, the intracellular pathways regulating fibroblast activation and matrix remodelling remain incompletely understood. Protein kinases are central regulators of cytoskeletal organisation, mechanotransduction and cell migration and therefore represent attractive therapeutic targets in fibrosis.

Protein kinase N2 (PKN2) is a Protein kinase C (PKC)-related serine/threonine kinase that regulates cytoskeletal organisation, cell adhesion and tissue morphogenesis. A recent genome-wide association study identified PKN2 as a determinant of forced vital capacity (FVC) decline in patients with IPF, suggesting a potential role in disease progression^20^. Supporting a potential role in lung biology, reduced PKN2 expression has been associated with defective epithelial apical junction formation, while studies in other systems have implicated PKN2 in fibroblast biology, embryonic development and tissue remodelling^20–25^. Together, these observations suggest that PKN2 may contribute to pulmonary fibrosis; however, its expression, regulation and functional role in IPF remains largely unknown.

Given the central contribution of fibroblast activation and tissue remodelling to fibrotic disease progression, understanding how PKN2 may regulate these processes could provide important mechanistic insights. Here, we investigate the expression, regulation and function of PKN2 in human pulmonary fibrosis and define its contribution to fibroblast activation and ECM remodelling.

## Methods

*Detailed experimental procedures are provided in the Supplementary Methods.*

### Human lung tissue and sputum samples

Human lung tissue and sputum samples were obtained from non-fibrotic control (NFC) donors and patients with ILD under approved ethical approvals (10/H0402/12, 22/EM/0159 and 07/MRE08/42). Non-fibrotic lung tissue was obtained from macroscopically normal regions of lungs resected for carcinoma at Glenfield Hospital, Leicester, UK. ILD and IPF lung tissue was obtained from diagnostic biopsies, explanted lungs or post-mortem tissue. Patient demographics are summarised in Supplementary Tables 1–4.

### Immunohistochemistry

Paraffin-embedded lung sections were stained for PKN2 using a rabbit monoclonal anti-PKN2 antibody (Thermo Fisher Scientific, #MA5-15877), antibody validation is described in supplementary figure 1 and ^26^. Images were acquired using brightfield microscopy and quantified using QuPath (v0.2.0). Detailed staining procedures are provided in the Supplementary Methods.

### Primary human cell isolation and culture

Primary airway and parenchymal fibroblasts were isolated from non-fibrotic and IPF lungs as previously described and cultured in Dulbecco’s Modified Eagle Medium supplemented with 10% fetal bovine serum^27–29^. Human airway epithelial cells, airway smooth muscle cells and fibroblasts used for comparative expression analyses were isolated and characterised as previously described^30^. All experiments were performed using cells at early passage (≤7). Mycoplasma-free status was confirmed before experimentation. Full cell isolation, characterisation and culture conditions are described in the Supplementary Methods.

### Cell stimulation and gene silencing

Fibroblasts were serum deprived prior to stimulation with recombinant human TGF-β1 where indicated. PKN2 knockdown was achieved using SMARTpool siRNA (Dharmacon) with a non-targeting siRNA control using INTERFERin® transfection reagent. Experimental conditions are described in the Supplementary Methods.

### RNA isolation and quantitative PCR

Total RNA was isolated from lung tissue, cultured cells and sputum using commercially available kits (Qiagen and Thermo Fisher Scientific) according to the manufacturers’ instructions. Gene expression was quantified by qRT-PCR using SYBR Green chemistry. Primer sequences and reaction conditions are provided in the Supplementary Methods.

### Cell viability

Cell viability following siRNA-mediated knockdown was assessed by trypan blue exclusion. Detailed protocols are provided in the Supplementary Methods.

### Enzyme linked immunosorbent assay

Secreted human procollagen Iα1, human vascular endothelial growth factor (VEGF) and human matrix metalloproteinase-1 (MMP-1) were quantified in fibroblast culture supernatants using commercially available ELISA kits according to the manufacturers’ instructions. Details are provided in the Supplementary Methods.

### Scratch wound assay

Isolated fibroblasts from NFC and IPF donors were transfected with siRNA as described above. After 24-hours of transfection, a scratch wound assay was performed and analysed as previously described^31^.

### Spatial Transcriptomic Reanalysis of Public Visium Data

Publicly available 10x Genomics Visium spatial transcriptomic data were obtained from Zenodo (doi: 10.5281/zenodo.10012934) and reanalysed to assess the *in situ* expression of *PKN2* using Scanpy^32^. Seven sections were processed including four healthy control and three IPF sections using the original celltype and niche annotation^33^.

Mean spot-level *PKN2* expression were compared between healthy and IPF sections and assessed using a two-sided Mann-Whitney U test, with each slide treated as the biological repeat.

### Quantitative real-time polymerase chain reaction (qRT-PCR)

cDNA was synthesised from ex vivo lung tissue, isolated human fibroblast and sputum RNA using SuperScript IV VILO Master Mix (ThermoFisher Scientific) or LunaScript® RT SuperMix Kit (New England Biolabs) according to manufacturing instructions using the ProFlex PCR System (Applied Biosystems, Life Technologies). qRT-PCR was performed using gene specific TaqMan primers to measure expression of PPIA (ThermoFisher Scientific, #Hs99999904_m1) and PKN2 (ThermoFisher Scientific, #Hs00178944_m1). qRT-PCR was carried out using a QuantStudio 5 Flex Real-Time PCR System (ThermoFisher Scientific).

### RNASeq

RNA sequencing was performed on NFC fibroblasts (n = 5) transfected with PKN2 or non-targeting control siRNA. Samples underwent quality control checks, library prep and were sequenced using Illumina PE150 technology. Sequencing was performed externally by Novogene (Novogene UK Company Limited, Cambridge, United Kingdom).

### Liquid chromatography mass spectrometry (LCMS)

Protein lysates from NFC fibroblasts (n = 3) following siRNA transfection with either PKN2 or non-targeting control siRNA were analysed by bottom-up quantitative LC-MS/MS using SP3 sample preparation and an Evosep One coupled to a timsTOF HT mass spectrometer. Raw data were processed using Spectronaut and ProteoScape. Detailed sample preparation, acquisition parameters and data processing are described in the Supplementary Methods.

### DNA methylation analysis

Genome-wide DNA methylation profiling was performed on cultured airway and lung parenchymal fibroblasts from non-fibrotic controls (airway, n = 8; parenchymal, n = 14) and patients with IPF (airway, n = 8; parenchymal, n = 8) using the Illumina HumanMethylationEPIC BeadChip array. Genomic DNA was isolated using the AllPrep DNA/RNA Mini Kit (Qiagen), bisulphite converted (EZ DNA Methylation Kit, Zymo Research), and hybridised to HumanMethylationEPIC BeadChips according to the manufacturers’ instructions. Raw IDAT files were processed in R using the minfi package with functional normalisation, probe quality filtering and batch correction. Differential methylation of CpG sites annotated to PKN2 was assessed using linear modelling with limma. Full details are provided in the Supplementary Methods.

### Gene expression microarray

Total RNA was isolated from matched fibroblast cultures using the AllPrep DNA/RNA Mini Kit (Qiagen) and analysed using the Affymetrix Human Gene 2.1 ST Array according to the manufacturer’s protocol. Raw CEL files were processed in R using GCRMA normalisation and batch correction prior to extraction of PKN2 expression data. Full details are provided in the Supplementary Methods.

### DNMT inhibition

Primary human lung fibroblasts were treated with increasing concentrations of the DNA methyltransferase inhibitor 5-aza-2′-deoxycytidine (5-aza; Sigma-Aldrich) for 96 hours under standard culture conditions. RNA and genomic DNA were isolated using the AllPrep DNA/RNA Mini Kit (Qiagen), and PKN2 mRNA expression was quantified by RT-qPCR using SYBR Green chemistry. Full experimental details, including treatment conditions and primer sequences, are provided in the Supplementary Methods.

### Zebrafish

All zebrafish experiments were performed in accordance with the UK Animals (Scientific Procedures) Act 1986 and approved by the University of Bristol Animal Welfare and Ethical Review Body. Wild-type and transgenic zebrafish lines were maintained under standard conditions as described previously^34,35^. Needle-stick wounds were performed in 4 days post-fertilisation (dpf) larvae, followed by treatment with the dual, selective PKN1/2 inhibitor PKN1/2-IN-1 (2 μM) or vehicle control. Wound healing, keratinocyte behaviour and collagen deposition were assessed by confocal microscopy and quantified using Fiji and custom image analysis pipelines. Full details of zebrafish husbandry, transgenic lines, wounding procedures, pharmacological treatment, imaging, image analysis and statistical methods are provided in the Supplementary Methods.

### Statistical Methods

#### General statistical analysis

GraphPad Prism (Version 9 and 10, GraphPad Software, San Diego, California, USA) was used for all data analysis (excluding RNA-sequencing, LC-MS proteomics and Zebrafish work). Data distribution and normality was assessed using the Shapiro-Wilk test, and appropriate parametric or non-parametric statistical tests used as stated in the figure legends. Statistical significance was defined as P < 0.05.

#### RNA sequencing and Proteomics Statistical Analysis

Differential expression analysis was performed in R using limma following data preprocessing, imputation and normalisation. Functional enrichment was performed using Metascape and STRING^36,37^. Drug repurposing analyses were performed using NeDRex within Cytoscape^38^. Full bioinformatic workflows are described in the Supplementary Methods.

## Results

### Loss of PKN2 is a feature of fibrotic lung remodelling

Immunohistochemical staining of paraffin-embedded lung tissue from non-fibrotic control (NFC: n = 12) and ILD (n = 6) donors demonstrated widespread PKN2 expression throughout the airway epithelium and lung parenchyma (Fig. 1A). Antibody specificity was confirmed by PKN2 knockdown in A549 epithelial cells and primary NFC fibroblasts (Supplementary Fig. 1 A and B, Figure 4A). Quantitative image analysis demonstrated significantly reduced PKN2 staining in ILD lung tissue compared with NFC (Mann-Whitney test, P = 0.0320; Fig. 1B), indicating that loss of PKN2 is a feature of fibrotic lung remodelling.

**Figure 1.**
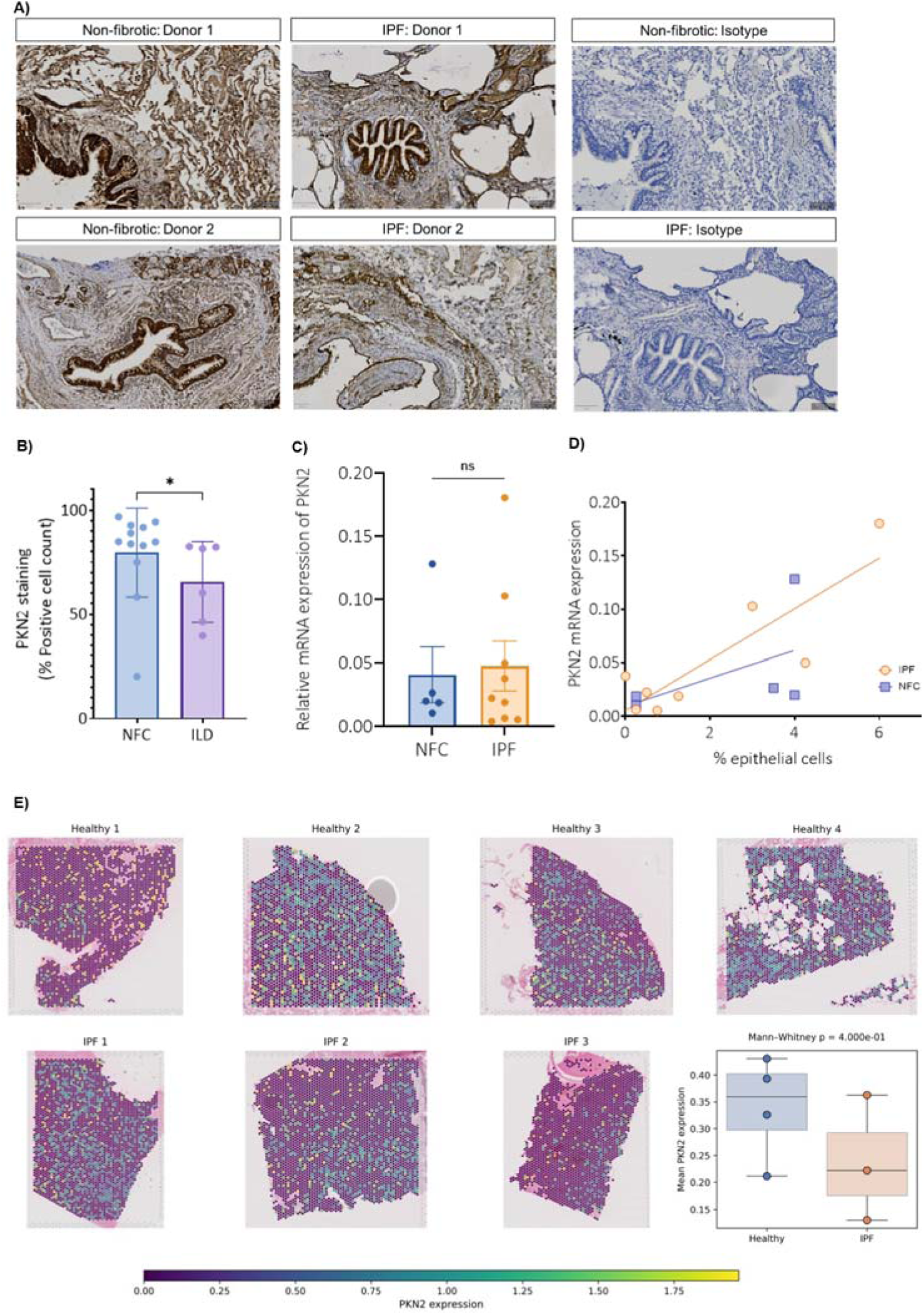
PKN2 expression is reduced in ILD lung tissue and is associated with epithelial cell abundance in induced sputum. A) Representative immunohistochemical staining of PKN2 in paraffin-embedded lung tissue sections from non-fibrotic control (NFC; n = 12) and interstitial lung disease (ILD; n = 6) donors. Representative isotype control staining is shown, scale bars 100μm. B) Quantification of PKN2 immunohistochemical staining demonstrating a reduced percentage of PKN2-positive cells in IPF compared with NFC lung tissue (Mann-Whitney test, *P = 0.0320). C) Relative PKN2 mRNA expression in induced sputum from NFC (n = 5) and IPF (n = 9) donors measured by qRT-PCR, Mann-Whitney test, P = ns. D) Correlation between sputum PKN2 mRNA expression and epithelial cell abundance in induced sputum from NFC (r^2^ = 0.2737, P = 0.365) and IPF donors (r^2^ = 0.7411, P = 0.006). E) Representative 10x Genomics Visium spatial transcriptomic maps showing PKN2 expression in healthy (n = 4) and IPF (n = 3) lung tissue. Box plot shows mean spot-level PKN2 expression for each donor (Mann-Whitney test, P= 0.40). Colour scale indicates normalised PKN2 expression.

To investigate whether PKN2 could be detected in a clinically accessible sample, we measured PKN2 mRNA in induced sputum from NFC (n = 5) and IPF (n = 9) donors. *PKN2* transcripts were readily detectable but did not differ significantly between disease groups (Fig. 1C). However, PKN2 expression positively correlated with the proportion of epithelial cells present within sputum samples (Fig. 1D), suggesting that sputum *PKN2* expression is influenced by epithelial cell abundance. No significant correlations were observed between *PKN2* expression, FVC or the proportions of neutrophils, macrophages, eosinophils or lymphocytes (Supplementary Fig. 1D), supporting epithelial cells as the predominant source of sputum *PKN2*.

Publicly available 10x Genomics Visium spatial transcriptomic datasets from healthy (n = 4) and IPF (n = 3) lungs were subsequently analysed^33^. *PKN2* expression was detected throughout both healthy and IPF lung tissue across multiple spatial regions (Fig. 1E). Mean spot-level *PKN2* expression was lower in IPF lungs than healthy controls, although this difference did not reach statistical significance (Mann–Whitney test, P = 0.40; Fig. 1E). Spatial transcriptomic analysis demonstrated widespread PKN2 expression with a trend towards lower expression in IPF tissue.

### Cell-specific analyses identify fibroblasts as a disease-relevant PKN2-expressing population

Analysis of publicly available GTEx lung cell expression data single-cell RNA sequencing datasets demonstrated that *PKN2* is broadly expressed across multiple pulmonary cell populations, including epithelial cells and fibroblasts (Fig. 2A)^39^.

**Figure 2.**
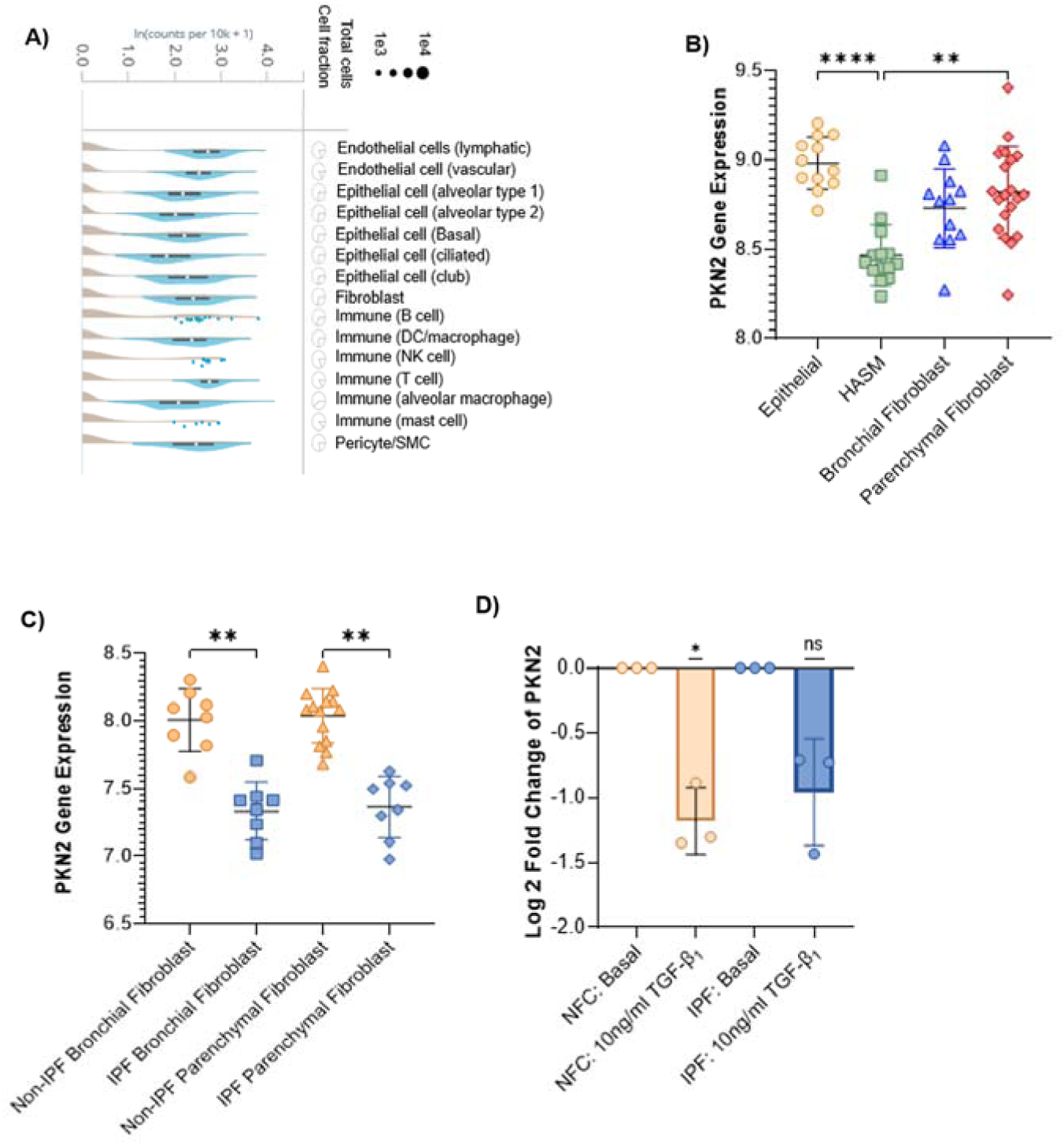
PKN2 is enriched in epithelial cells and fibroblasts, reduced in IPF fibroblasts, and downregulated by TGF-β1. A) Publicly available GTEx lung cell expression data demonstrating PKN2 expression across multiple pulmonary cell types. B) Affymetrix Human Gene 2.1 ST microarray analysis demonstrating PKN2 expression in primary human bronchial epithelial cells (n = 21), airway smooth muscle cells (HASM; n = 9), bronchial fibroblasts (n = 18) and parenchymal fibroblasts (n = 24). PKN2 expression was significantly lower in HASM than bronchial epithelial cells (P < 0.0001) and parenchymal fibroblasts (P= 0.0022). Statistical analysis was performed using a Kruskal-Wallis test with multiple comparisons. C) Affymetrix Human Gene 2.1 ST microarray analysis of primary fibroblasts demonstrating significantly reduced PKN2 expression in IPF bronchial fibroblasts (Non-IPF, n = 8; IPF, n = 8; P = 0.0044) and IPF parenchymal fibroblasts (Non-IPF, n = 14; IPF, n = 8; P = 0.0011). Statistical analysis was performed using a Kruskal-Wallis test with multiple comparisons. D) PKN2 mRNA expression following TGF-β1 stimulation (10 ng/ml) in primary human lung fibroblasts. TGF-β1 significantly reduced PKN2 expression in non-fibrotic control (NFC) fibroblasts (n = 3) and produced a similar, non-significant reduction in IPF fibroblasts (n = 3).

To determine whether fibroblasts represent a disease-relevant PKN2-expressing population, we interrogated independent Affymetrix gene expression datasets from isolated primary human bronchial epithelial cells (n = 12), airway smooth muscle (HASM; n = 15), bronchial fibroblasts (n = 13) and parenchymal fibroblasts (n = 20). *PKN2* expression was readily detected in all cell types, although expression was significantly lower in HASM than bronchial epithelial cells or parenchymal fibroblasts (Kruskal–Wallis, P < 0.0001 and P = 0.0022, respectively) (Fig. 2C).

We next examined whether *PKN2* mRNA expression was altered in disease-derived cultured fibroblasts. *PKN2* mRNA expression was significantly reduced in IPF bronchial fibroblasts (n = 8) compared with non-IPF bronchial fibroblast controls (n = 8) and was similarly reduced in IPF parenchymal fibroblasts (n = 8) compared with non-IPF parenchymal fibroblasts (n = 14) (Fig. 2D).

Because, transforming growth factor-β1 (TGF-β1) is a central driver of fibroblast activation in pulmonary fibrosis, we examined its effects on *PKN2* expression in isolated primary human lung fibroblasts. Dose-response experiments identified that TGF-β1 suppressed *PKN2* mRNA expression, with maximal suppression at 10 ng/ml TGF-β1 (Supplementary Fig. 1D). This downregulation was sustained over 72 h (Supplementary Fig. 1E). *PKN2* expression was significantly reduced following 24 h TGF-β1 10 ng/ml stimulation in NFC fibroblasts (P = 0.0154), with a similar trend observed in IPF fibroblasts (Fig. 2E).

Collectively, these findings identify fibroblasts as a disease-relevant PKN2-expressing cell population in which *PKN2* expression is consistently reduced in IPF and further suppressed by profibrotic TGF-β1 signalling.

### PKN2 exhibits altered DNA methylation in IPF-derived airway and parenchymal fibroblasts

Given the consistent reduction in *PKN2* expression in IPF-derived fibroblasts, we next examined whether altered DNA methylation contributed to its dysregulation. DNA methylation was interrogated across the entire *PKN2* locus. Thirty annotated CpG sites passed quality control and were analysed by linear regression in airway and parenchymal fibroblasts from non-IPF and IPF donors.

Four CpG sites were differentially methylated positions (DMPs) in airway fibroblasts and seven in parenchymal fibroblasts (nominal P<0.005; Fig. 3A,C). cg25096094 was differentially methylated in both fibroblast populations. Most DMPs mapped to gene body or OpenSea regions, whereas cg13879769 was located within the south shore of a CpG island. To determine whether these methylation changes were associated with gene expression, we correlated DNA methylation β-values with *PKN2* mRNA expression. Significant associations were identified for multiple CpG sites in both airway and parenchymal fibroblasts, with both positive and negative correlations depending on genomic location (Fig. 3B,D), suggesting locus-specific epigenetic regulation of *PKN2* expression. To further investigate the relationship between DNA methylation and *PKN2* expression, fibroblasts were treated with the DNA methyltransferase inhibitor 5-aza-2′-deoxycytidine. DNA methyltransferase (DNMT) inhibition significantly altered *PKN2* expression in non-fibrotic fibroblasts, whereas only a modest effect was observed in IPF fibroblasts (P = 0.05) (Fig. 3E,F). Collectively, these findings identify *PKN2* as an epigenetically dysregulated gene in IPF fibroblasts and suggest that altered DNA methylation contributes to the regulation of PKN2 expression in disease.

**Figure 3.**
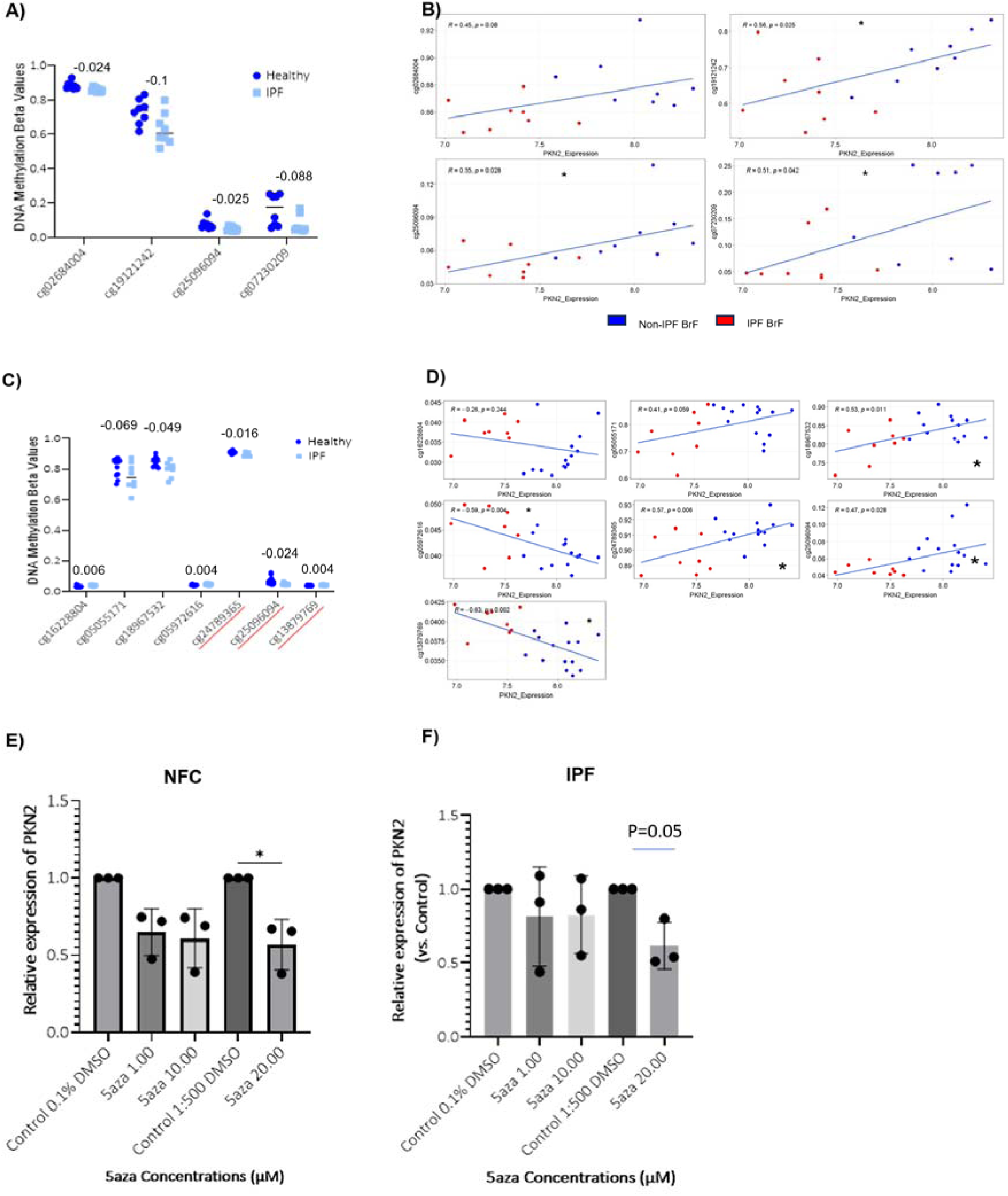
PKN2 exhibits differential DNA methylation in IPF-derived airway and parenchymal fibroblasts. (A) DNA methylation of CpG sites annotated to PKN2 in airway fibroblasts (BrF) from non-fibrotic control (NFC) and IPF donors measured using the Illumina HumanMethylationEPIC BeadChip array. Differentially methylated CpG sites identified by linear regression are shown. (B) Correlation analysis between PKN2 mRNA expression and DNA methylation (β-values) at differentially methylated CpG sites in airway fibroblasts. (C) DNA methylation of CpG sites annotated to PKN2 in lung parenchymal fibroblasts (LgF) from NFC and IPF donors measured using the Illumina HumanMethylationEPIC BeadChip array. Differentially methylated CpG sites identified by linear regression are shown. CpG sites underlined in red remained significant following Benjamini–Hochberg false discovery rate (FDR) correction (P < 0.05). (D) Correlation analysis between PKN2 mRNA expression and DNA methylation (β-values) at differentially methylated CpG sites in lung parenchymal fibroblasts. (E,F) Effect of the DNA methyltransferase inhibitor 5-aza-2′-deoxycytidine (5-aza) on PKN2 mRNA expression in primary NFC (E) and IPF (F) fibroblasts (n = 3). Relative PKN2 expression was determined by RT-qPCR following treatment with increasing concentrations of 5-aza. Bars represent mean ± SEM. Statistical significance is indicated where appropriate.

### Loss of PKN2 promotes ECM remodelling and fibroblast activation

Having established that *PKN2* expression was reduced in ILD lung tissue and fibroblasts, and further suppressed by TGF-β1, we next investigated the consequences of *PKN2* depletion on the fibroblast transcriptome. *PKN2* knockdown was confirmed in NFC fibroblasts using qRT-PCR (n = 7) (Fig. 4A). Knockdown was optimised using pooled siRNA, resulting in efficient gene silencing without affecting cell viability.

**Figure 4.**
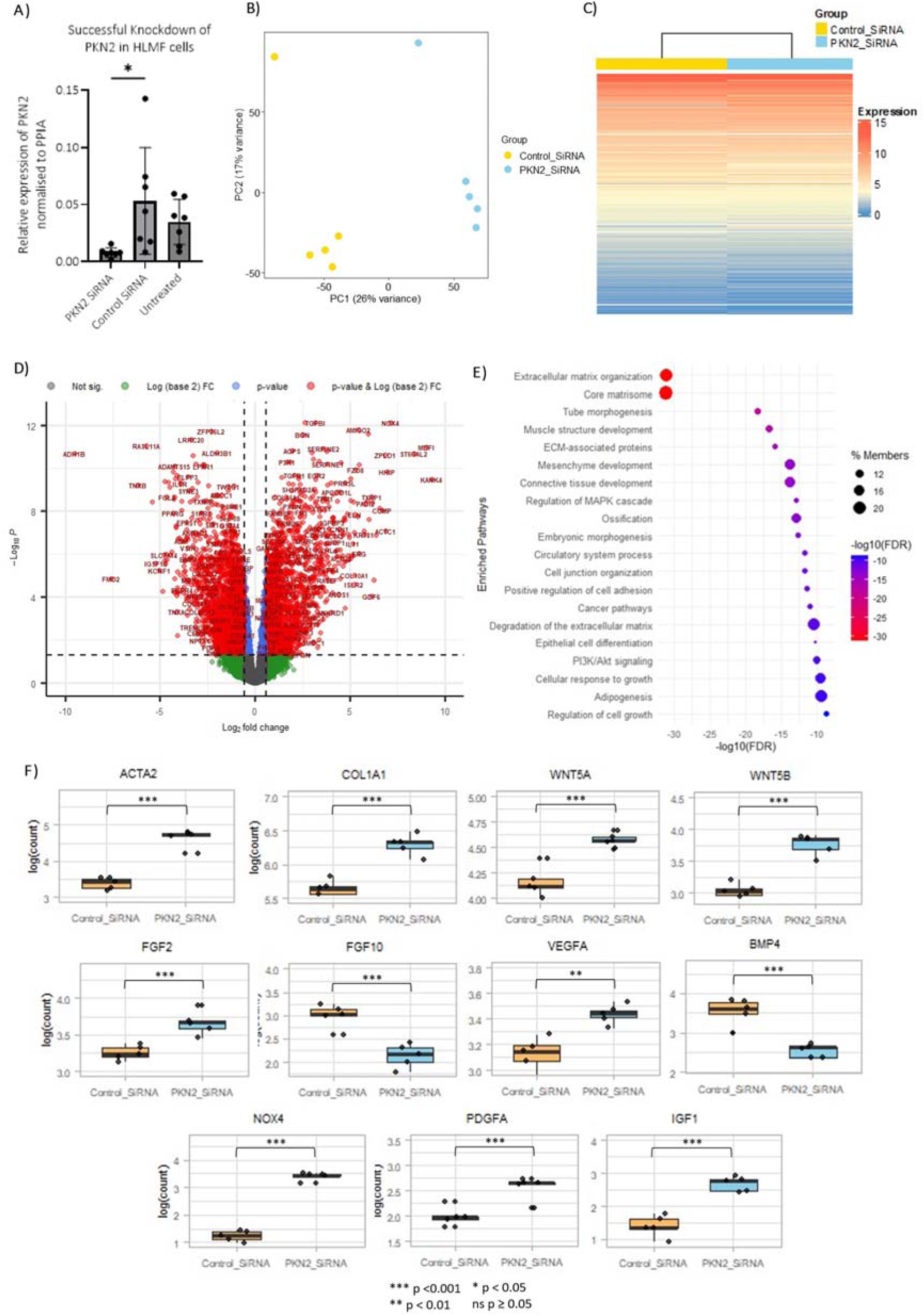
PKN2 knockdown reprogrammes the fibroblast transcriptome towards a profibrotic state. RNA sequencing was performed on non-fibrotic control (NFC) fibroblasts following PKN2 or non-targeting control siRNA transfection (n = 5 biological donors). (A) qRT-PCR confirmation of efficient PKN2 knockdown following siRNA transfection in non-fibrotic control (NFC) fibroblasts (n = 7). PKN2 expression was normalised to PPIA. (B) Principal component analysis (PCA) demonstrating separation of myofibroblasts according to siRNA treatment. (C) Unsupervised hierarchical clustering and heatmap of differentially expressed genes following PKN2 knockdown. (D) Volcano plot showing differentially expressed genes following PKN2 knockdown compared with non-targeting control siRNA. Differentially expressed genes were identified using an adjusted P < 0.05 and |log₂ fold change| > 0.58. (E) Gene ontology pathway enrichment analysis highlighting the top enriched biological pathways following PKN2 knockdown. (F) Expression of selected differentially expressed genes associated with extracellular matrix remodelling, fibroblast activation and profibrotic signalling following PKN2 knockdown. Boxes represent the median and interquartile range, whiskers indicate the minimum and maximum values, and individual points represent biological replicates. Statistical significance is indicated as *P < 0.05, **P < 0.01, and ***P < 0.001, n = 5.

Bulk RNA sequencing was performed on NFC fibroblasts transfected with either non-targeting control or *PKN2* siRNA (n = 5). Principal component analysis demonstrated clear segregation of non-targeting control siRNA and *PKN2*-depleted fibroblasts along PC1 (26% variance), indicating distinct transcriptional profiles (Fig. 4B). Unsupervised hierarchical clustering similarly separated samples according to treatment (Fig. 4C), confirming that loss of *PKN2* induces widespread transcriptional remodelling.

Differential expression analysis identified 2,240 upregulated and 2,769 downregulated genes following *PKN2* knockdown (adjusted P < 0.05, log2(FC) > 0.58; Fig. 4D, Supplementary Table 5). Gene ontology analysis demonstrated significant enrichment of pathways associated with ECM organisation, core matrisome components, ECM-associated proteins, cell adhesion, cell junction organisation and MAPK signalling (Fig. 4E). These pathways are characteristic of profibrotic transcriptional programmes previously identified in IPF lung tissue, primary fibroblasts and TGFβ1-stimulated human lung tissue^40,41^.

To further define this phenotype, we examined individual genes from RNAseq data with established roles in fibroblast activation and matrix remodelling (Fig. 4F). COL1A1 and ACTA2, key markers of ECM deposition and myofibroblast differentiation, were significantly upregulated following PKN2 knockdown, both ***P < 0.001, n = 5^42,43^. Expression of the non-canonical WNT ligands WNT5A and WNT5B, which regulate fibroblast activation and tissue remodelling, was also increased, ***P < 0.001, ^44,45^. In addition, PDGFA, IGF1, VEGFA, FGF2 and NOX4, all implicated in fibroblast proliferation, angiogenesis or oxidative stress, were significantly upregulated, whereas the reparative growth factor FGF10 and the antifibrotic mediator BMP4 were significantly reduced, all P < 0.01^46–49^.

Collectively, these findings indicate that loss of *PKN2* reprogrammes fibroblasts towards a profibrotic state, characterised by ECM remodelling and fibroblast activation. We therefore investigated whether these transcriptional alterations were recapitulated at the protein level.

### Loss of PKN2 induces coordinated profibrotic remodelling of the fibroblast proteome

To determine whether the transcriptional changes induced by PKN2 depletion were reflected at the protein level, we performed untargeted quantitative LC-MS/MS on NFC-derived fibroblasts transfected with either non-targeting control or PKN2 siRNA (n = 3).

Principal component analysis demonstrated separation of non-targeting control and PKN2-depleted fibroblasts based on their proteomic profiles along PC2 (10% variance)(Fig. 5A), indicating that PKN2 knockdown induces measurable, albeit more modest, changes at the protein level than observed by RNA sequencing. Differential expression analysis identified 392 upregulated and 324 downregulated proteins (adjusted P < 0.05), with 64 proteins remaining significantly altered following application of a log2(FC) > 0.58 fold change threshold (Fig. 4B, Supplementary Table 6).

**Figure 5.**
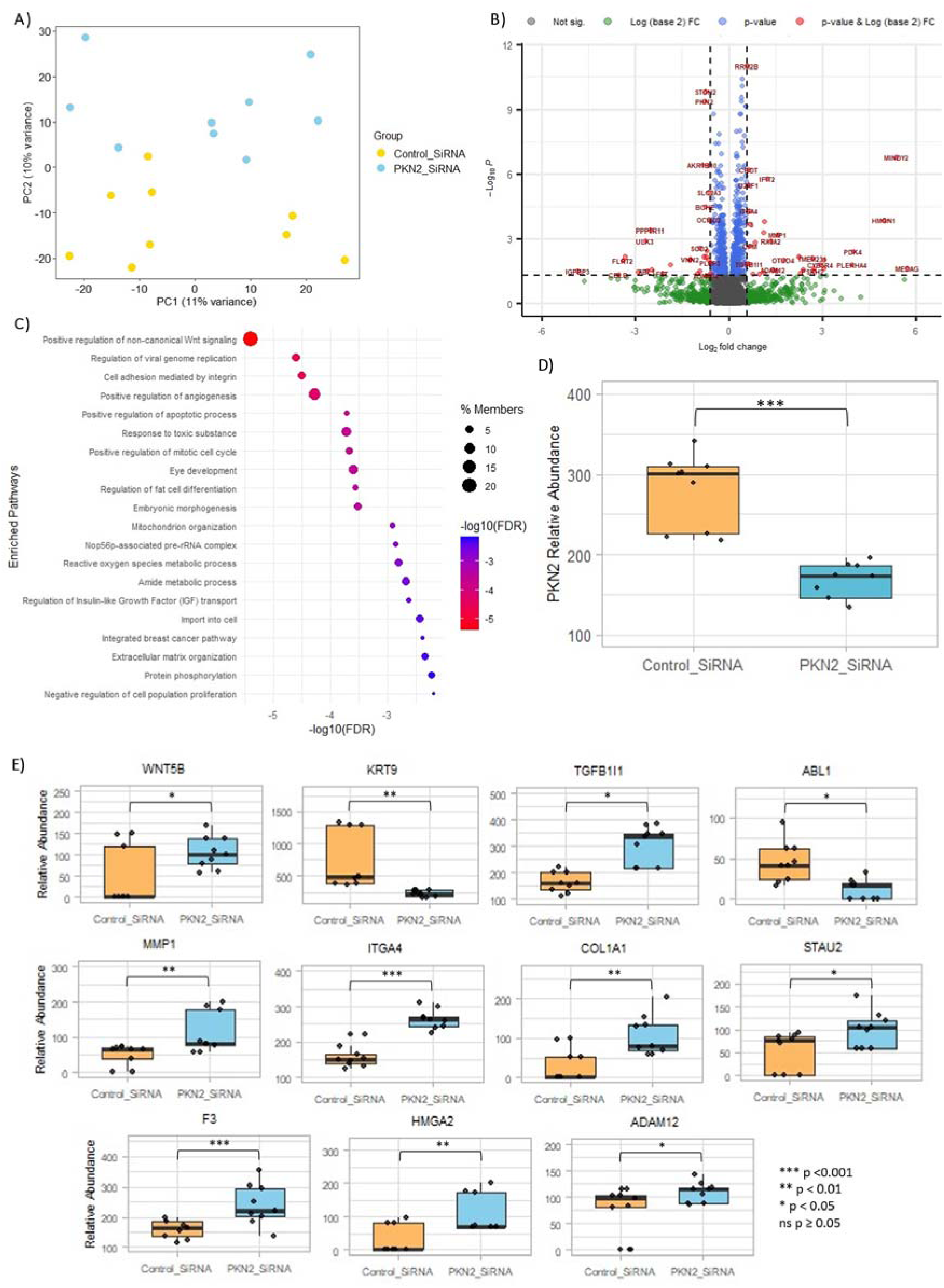
Quantitative proteomic profiling identifies PKN2-dependent remodelling of the fibroblast proteome. **(A)** Principal component analysis (PCA) of LC-MS/MS proteomic profiles from non-fibrotic control (NFC) fibroblasts transfected with non-targeting control or PKN2 siRNA (n = 3 biological donors). Samples are coloured according to treatment group. **(B)** Volcano plot showing differentially expressed proteins following PKN2 knockdown compared with non-targeting control siRNA. Differentially expressed proteins were identified using an adjusted p < 0.05 and |log₂ fold change| > 0.58. **(C)** Gene ontology pathway enrichment analysis of significantly dysregulated proteins following PKN2 knockdown, highlighting the top enriched biological pathways. **(D)** Relative PKN2 protein abundance measured by LC-MS/MS following non-targeting control or PKN2 siRNA transfection. **(E)** Relative abundance of selected significantly dysregulated proteins associated with extracellular matrix remodelling, cell adhesion and profibrotic signalling following PKN2 knockdown. Boxes represent the median and interquartile range, whiskers indicate the minimum and maximum values, and individual points represent biological replicates. Statistical significance is indicated as *p < 0.05, **p < 0.01, and ***p < 0.001.

Pathway enrichment analysis revealed significant enrichment of biological processes associated with non-canonical WNT signalling, integrin-mediated cell adhesion, ECM organisation, reactive oxygen species metabolism and angiogenesis (Fig. 5C), closely mirroring the transcriptional programmes identified by RNA sequencing.

Interestingly, PKN2 siRNA knockdown resulted in marked reduction in PKN2 protein abundance (Fig. 5D). Examination of individual proteins demonstrated increased abundance of several implicated in fibroblast activation and ECM remodelling (Fig. 5E). These included COL1A1, WNT5B, ITGA4, TGFB1I1, ADAM12 and HMGA2, together with increased MMP1, consistent with active ECM turnover.

Collectively, these findings demonstrate that loss of PKN2 induces coordinated transcriptional and proteomic remodelling, converging on ECM organisation, cell adhesion and non-canonical WNT signalling.

### Multi-omics integration reveals coordinated PKN2-dependent regulation of ECM and profibrotic signalling

To define the relationship between transcriptional and proteomic responses to PKN2 depletion, we integrated the RNA sequencing and quantitative proteomic datasets. This approach enabled identification of molecular changes that were conserved across both expression layers, highlighting biologically robust consequences of PKN2 loss.

Comparison of RNA and protein expression across the 5,563 shared features demonstrated an overall positive relationship between the two datasets, with approximately 60% of molecules showing concordant expression changes (Fig. 6A). This level of concordance is consistent with previous studies comparing transcriptomic and proteomic datasets, reflecting both coordinated regulation and post-transcriptional control^50^. PKN2 itself demonstrated concordant reductions at both the transcript and protein level, confirming effective target depletion (Fig. 4A & 5D).

**Figure 6.**
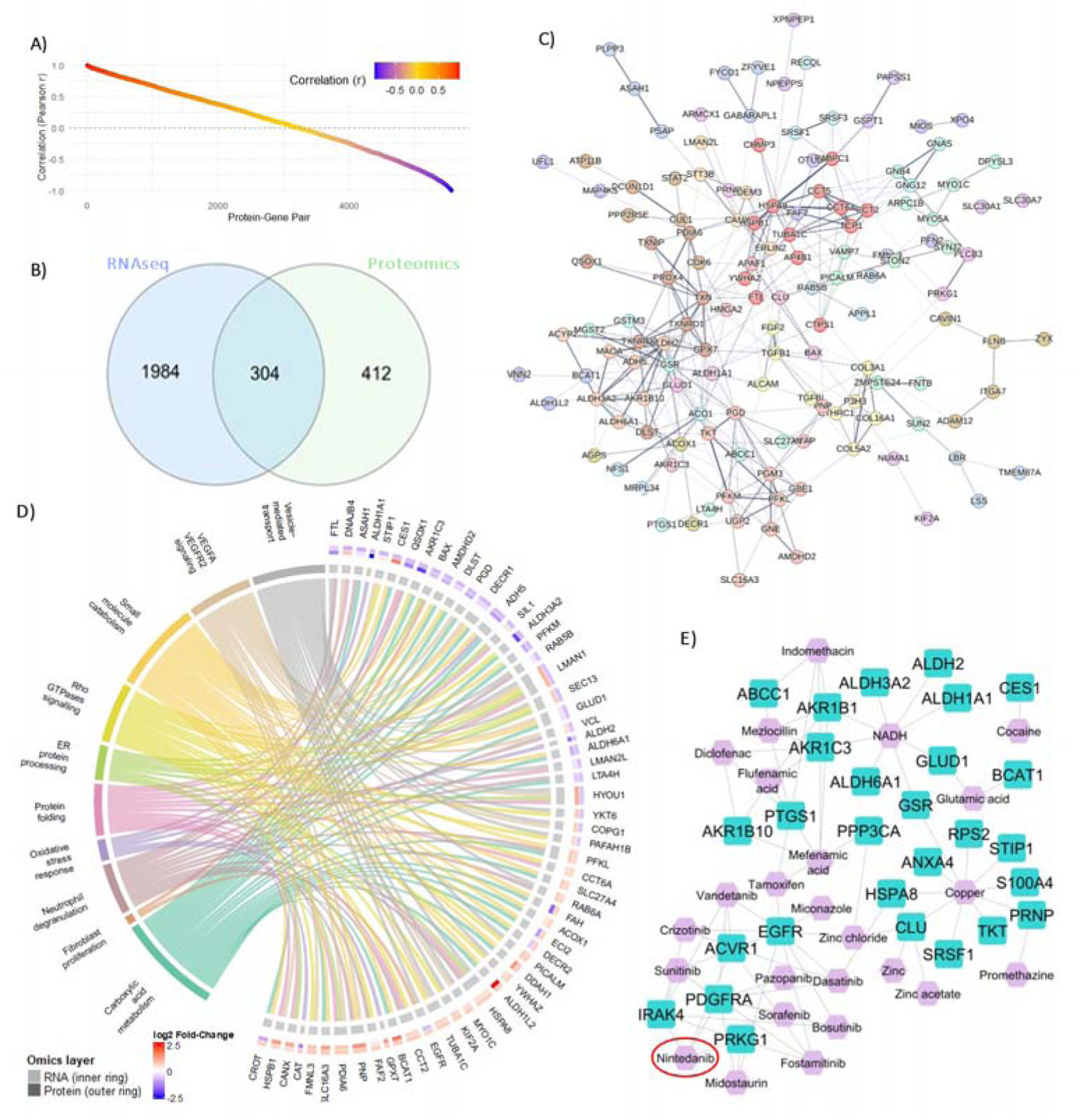
Integrated transcriptomic and proteomic analyses identify conserved PKN2-regulated molecular networks. **(A)** Correlation analysis comparing transcriptomic (RNA-seq) and proteomic (LC-MS/MS) expression changes across the 5,563 genes detected in both datasets following PKN2 knockdown in non-fibrotic control (NFC) fibroblasts. Each point represents a matched transcript–protein pair, coloured according to the Pearson correlation coefficient. **(B)** Venn diagram showing the overlap between significantly differentially expressed transcripts and proteins following PKN2 knockdown. A total of 304 features were significantly altered in both datasets. **(C)** STRING protein-protein interaction network of the 304 shared differentially expressed features, illustrating highly interconnected molecular networks regulated by PKN2. Nodes represent proteins and edges represent experimentally validated or predicted protein-protein interactions. **(D)** Integrated pathway analysis of concordantly dysregulated transcriptomic and proteomic features. Chord diagram illustrating the association between significantly enriched biological pathways and overlapping differentially expressed genes/proteins. Inner ring indicates RNA-seq log₂ fold change and outer ring indicates protein log₂ fold change. **(E)** Network-based drug repurposing analysis of the integrated multi-omics signature using NeDRex. The top 25 predicted candidates targeting the PKN2-regulated network ranked using the TrustRank algorithm are shown in blue and candidate compounds in purple. Nintedanib, an approved antifibrotic therapy for idiopathic pulmonary fibrosis, was identified among the highest-ranked candidate compounds.

Among the differentially expressed molecule common to both datasets, 304 were significantly altered at both the transcriptomic and proteomic levels (Fig. 6B), with more than half (56%,169/304 proteins) exhibiting concordant directionality of change (Supplementary Table 7). These findings indicate that a substantial proportion of the molecular response to PKN2 depletion is maintained from transcription through to protein abundance.

Protein-protein interaction network analysis of these shared molecules identified highly interconnected modules enriched for ECM organisation, collagen biology, protein folding and redox regulation (Fig. 6C), demonstrating that PKN2 depletion remodels coordinated biological networks rather than isolated molecular pathways.

Integrated pathway analysis of the 304 significantly altered features at both the transcriptomic and proteomic level further highlighted enrichment of fibroblast proliferation, VEGFA–VEGFR2 signalling, RHO GTPase signalling and cellular metabolic pathways (Fig. 6D). Several key regulators, including YWHAZ, MYOC1 and CANX, were concordantly upregulated at both transcript and protein levels, whereas RAB5B and PFKM were consistently reduced, supporting coordinated regulation across multiple molecular layers.

Finally, network-based drug repurposing analysis based on the conserved differentially expressed features identified multiple candidate therapeutics predicted to target the PKN2-regulated molecular network (Fig. 6E). Notably, Nintedanib, an approved antifibrotic therapy for IPF, emerged among the highest-ranked candidates, providing independent support for the biological relevance of the PKN2-regulated network and suggesting that PKN2 loss converges on clinically actionable profibrotic pathways.

### Loss of PKN2 increases secretion of ECM remodelling mediators and promotes fibroblast wound repair

VEGF, pro-collagen Iα1 and MMP-1 are key mediators of ECM remodelling and tissue repair in IPF. VEGF regulates angiogenesis and fibroblast survival and is implicated in fibrotic progression, whereas MMP-1 is a collagenase involved in ECM turnover^51–53^. Type I collagen is the predominant fibrillar collagen deposited within the fibrotic lung^42,54^.

To determine whether transcriptional and proteomic changes induced by PKN2 depletion translated into altered protein secretion, conditioned media from fibroblasts transfected with non-targeting control or PKN2 siRNA were analysed by ELISA. PKN2 knockdown significantly increased VEGF secretion in both NFC (n = 7, P = 0.0146) and IPF (n = 7, P = 0.0038) fibroblasts (Fig. 7A). Similarly, total MMP-1 secretion was significantly increased following PKN2 knockdown in NFC (n = 6, P = 0.0212) and IPF (n = 6, P = 0.0449) fibroblasts (Fig. 7B). In contrast, secretion of pro-collagen Iα1 was unchanged in either NFC (n = 7, P > 0.9999) or IPF (n = 6, P = 0.3693) fibroblasts (Fig. 7C).

**Figure 7.**
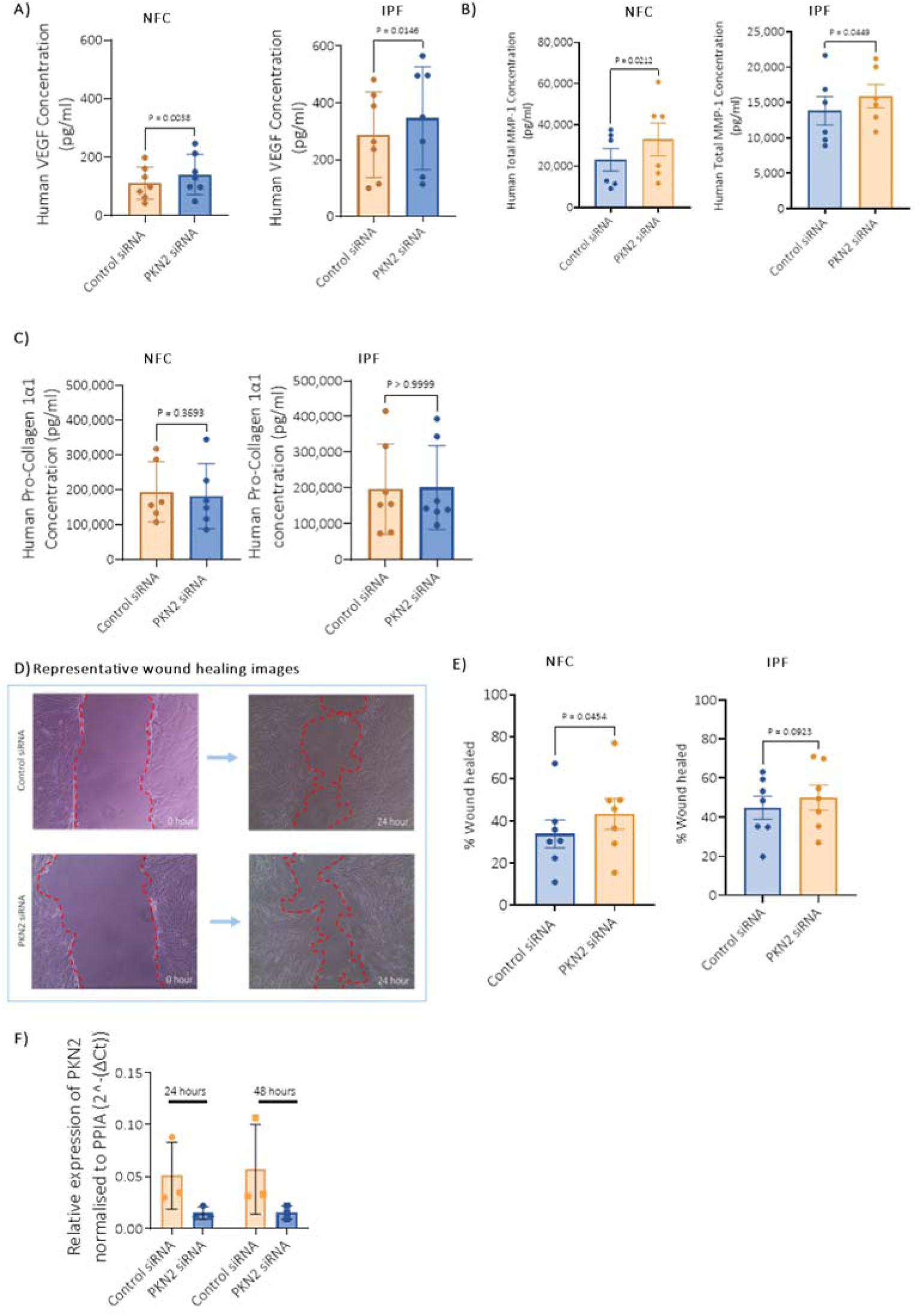
PKN2 knockdown increases VEGF and MMP-1 secretion and accelerates wound closure in fibroblasts. Primary human lung fibroblasts were transfected with pooled PKN2 siRNA or non-targeting control siRNA for 48 h before analysis. **(A)** VEGF secretion measured by ELISA was significantly increased following PKN2 knockdown in both non-fibrotic control (NFC; n = 7, P = 0.0146) and IPF fibroblasts (n = 7, p = 0.0038). **(B)** Pro-collagen Iα1 secretion was not significantly altered following PKN2 knockdown in either NFC (n = 7, P > 0.9999) or IPF fibroblasts (n = 6, P = 0.3693). **(C)** Total MMP-1 secretion measured by ELISA was significantly increased following PKN2 knockdown in both NFC (n = 6, P = 0.0212) and IPF myofibroblasts (n = 6, P = 0.0449). **(D)** Representative phase-contrast images of scratch wound assays performed in NFC fibroblasts immediately after wounding (0 h) and 24 h following treatment with non-targeting control or PKN2 siRNA. Wound margins are indicated by the red dashed lines. **(E)** Quantification of wound closure demonstrated significantly accelerated wound repair following PKN2 knockdown in NFC fibroblasts (n = 7, P = 0.0454), whereas IPF fibroblasts did not reach significance (n = 7, p = 0.0923). **(F)** qRT-PCR analysis confirmed sustained PKN2 knockdown at both 24 h and 48 h following wounding (NFC; n = 3). PKN2 expression was normalised to PPIA.

To investigate whether these molecular changes altered fibroblast behaviour, scratch wound assays were performed. PKN2 knockdown significantly accelerated wound closure in NFC fibroblasts after 24 h (n = 7, P = 0.0454), whereas IPF fibroblasts showed the same trend but did not reach significance (n = 7, P = 0.0923) (Fig. 7D,E). PKN2 knockdown was maintained throughout the assay, as confirmed by RT-qPCR (Fig. 7F).

Together, these findings demonstrate that loss of PKN2 promotes a functional phenotype characterised by enhanced wound closure and increased secretion of matrix-remodelling mediators, consistent with the transcriptomic and proteomic enrichment of ECM organisation, cell adhesion and WNT signalling pathways.

### PKN Inhibition disrupts pigmentation, re-epithelialisation, and collagen alignment in zebrafish wounds

Our results in scratch wound assays following PKN2 knockdown suggested that PKN2 could play a role in wound healing *in vivo*. To investigate this, we exploited the optical transparency of zebrafish larvae, an established model of wound healing^55,56^, together with transgenic reporters that enable live imaging of the wound epidermis and collagen remodelling ^56,57^. Needle-stick wounds were generated in the flank tissue of 4 days post-fertilisation (dpf) larvae above the cloaca, followed by treatment with the selective PKN inhibitor PKN1/2-IN-1 (PKNi; 2 μM) or vehicle (DMSO) (Fig. 8A).

**Figure 8.**
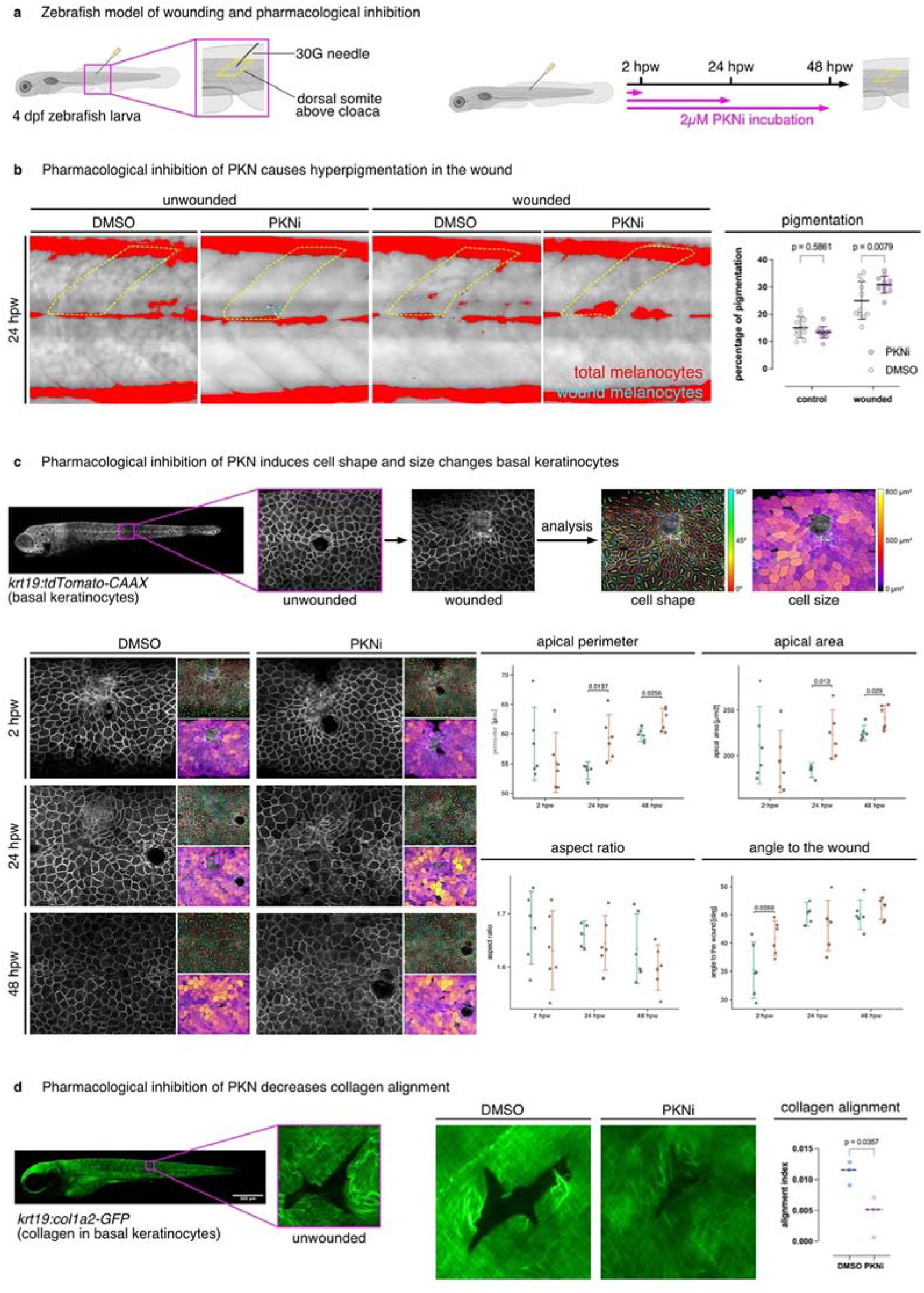
Pharmacological inhibition of PKN disrupts wound repair, epithelial organisation and collagen alignment in zebrafish larvae. A) Schematic of the zebrafish wound-healing model. Four days post-fertilisation (dpf) larvae were wounded in the dorsal somite above the cloaca using a 30G needle and incubated with 2 μM PKN1/2 inhibitor (PKNi) or vehicle (DMSO). Larvae were analysed at 2, 24 and 48 hours post-wounding (hpw). B) Representative images and quantification of wound hyperpigmentation at 24 hpw demonstrating increased melanocyte accumulation within wounds following PKN inhibition (n ≥ 10). C) Representative confocal images of wounded Tg(krt19:tdTomato-CAAX) larvae showing basal keratinocyte morphology following treatment with DMSO or PKNi. Morphometric analysis demonstrated increased keratinocyte apical perimeter and apical area at 24 and 48 hpw, while aspect ratio was unchanged. A transient increase in the angle of cell elongation relative to the wound was observed at 2 hpw (n = 6). D) Representative confocal images of wounded Tg(krt19:col1a2-GFP) larvae and quantification of collagen fibre alignment within the wound. Pharmacological inhibition of PKN significantly reduced the collagen alignment index, indicating impaired collagen remodelling during wound repair (n = 3).

Under normal conditions, needle stick wounds made to the flank tissue of the fish induce a transient hyperpigmentation response, where melanocytes and melanoblasts migrate to the wound ^58^. In zebrafish, we quantified the area of the wounded somite covered by melanocytes and found that by 24 hours post wounding (hpw) treatment with PKNi led to an increase in the pigmented area within the wounds (Figure 8B).

To determine whether PKN signalling regulates re-epithelialisation, we analysed wounded Tg(krt19:tdTomato-CAAX) larvae, in which basal keratinocyte membranes are fluorescently labelled^57^. Morphometric analysis revealed that PKNi significantly increased keratinocyte apical perimeter and apical area at 24 and 48 hpw, whereas cell aspect ratio remained unchanged. In contrast, the angle of cell elongation relative to the wound was transiently increased at 2 hpw, indicating altered early epithelial organisation during wound closure (Fig. 8C).

Because coordinated epithelial repair is closely linked to ECM remodelling, we next examined collagen organisation using Tg(krt19:col1a2-GFP) larvae, whose basal keratinocytes express GFP-tagged collagen 1a2^56^. As expected, needle injury initially disrupted the orthogonal collagen network, followed by progressive deposition and remodelling of collagen fibres during healing. Pharmacological inhibition of PKN significantly reduced the collagen alignment index, demonstrating impaired restoration of the organised collagen architecture within the wound (Fig. 8D).

Collectively, these findings demonstrate that inhibition of PKN signalling alters multiple stages of tissue repair *in vivo*, including epithelial organisation and ECM remodelling. Together with our transcriptomic, proteomic and functional analyses in primary human fibroblasts, these data identify PKN2 as a conserved regulator of wound repair and matrix organisation, supporting its potential role in the pathogenesis of pulmonary fibrosis.

## Discussion

IPF is characterised by persistent fibroblast activation and aberrant ECM remodelling, yet the molecular mechanisms governing these processes remain incompletely understood. Here, we identify PKN2 as a previously unrecognised regulator of human lung fibroblast homeostasis. We demonstrate that PKN2 expression is reduced in both fibrotic lung tissue and in both bronchial and parenchymal fibroblasts isolated from patients with ILD, and is further suppressed by the profibrotic cytokine TGF-β1. Loss of PKN2 promotes widespread molecular and functional changes consistent with fibroblast activation. By integrating epigenetic profiling, transcriptomics, proteomics and functional analyses, our study identifies PKN2 as a negative regulator of ECM remodelling and tissue repair, and provides a firm link to the previous GWAS signal identifying PKN2 as a potential driver of IPF.

The first key finding of this study is that reduced PKN2 expression is associated with altered DNA methylation. We identified multiple differentially methylated CpG sites across the PKN2 locus in both bronchial and parenchymal fibroblasts, including a shared CpG site present in both fibroblast populations, suggesting conserved epigenetic regulation. Although inhibition of DNA methyltransferases using 5-aza only partially restored PKN2 expression, these findings support a functional role for DNA methylation in regulating PKN2 transcription rather than representing the sole mechanism of repression. Epigenetic dysregulation is increasingly recognised as a key driver of fibroblast activation and regulation in respiratory conditions ^59,60^, and our data identify PKN2 as a novel fibrosis-associated gene that may be regulated, at least in part, through DNA methylation.

Transcriptomic profiling demonstrated that loss of PKN2 induced widespread transcriptional reprogramming of human lung fibroblasts. Gene ontology analysis identified enrichment of ECM organisation, core matrisome, cell adhesion, cell junction organisation and MAPK signalling pathways, together with increased expression of established profibrotic mediators including COL1A1, ACTA2, WNT5A, WNT5B, PDGFA, IGF1, NOX4 and VEGFA. The transcriptomic changes induced by PKN2 depletion closely resemble molecular signatures previously reported in IPF lung tissue and experimental models of pulmonary fibrosis, particularly enrichment of ECM organisation, cell adhesion and WNT signalling pathways^40,41^, supporting the biological relevance of the PKN2-dependent programme. These findings indicate that PKN2 functions as a negative regulator of fibroblast activation, with its loss shifting fibroblasts towards a transcriptional programme that favours ECM remodelling and tissue fibrosis.

Proteomic analysis demonstrated that these transcriptional changes were reflected at the protein level. Although fewer proteins were differentially expressed than transcripts, likely in part due to the smaller sample size, enrichment of non-canonical WNT signalling, ECM organisation, integrin-mediated adhesion and reactive oxygen species pathways closely mirrored the RNA sequencing data. Furthermore, proteins central to matrix remodelling, including COL1A1, MMP1, ADAM12, WNT5B and ITGA4, were significantly dysregulated, indicating that PKN2-dependent transcriptional changes are translated into biologically relevant alterations in the fibroblast proteome. Notably, these proteomic changes closely resemble those identified in our recently published quantitative proteomic analysis of a human *ex vivo* model of IPF, in which ECM remodelling, WNT signalling and oxidative stress pathways were similarly enriched^61^.

A major strength of this study is the integration of epigenetic, transcriptomic, proteomic and functional datasets, allowing us to define the consequences of PKN2 loss across multiple biological levels. Although induction of α-smooth muscle actin was not observed at the protein level, highlighting the imperfect relationship between transcript and protein abundance, integration of both datasets identified a conserved core of dysregulated profibrotic pathways. This systems-level approach provides greater confidence that loss of PKN2 drives coordinated remodelling of ECM organisation, cell adhesion, WNT signalling and angiogenic networks rather than isolated changes in individual genes.

An interesting finding was the differential regulation of collagen homeostasis. Although COL1A1 transcript and intracellular protein abundance increased following PKN2 depletion, secretion of procollagen Iα1 remained unchanged over the experimental time course. Collagen biosynthesis is tightly regulated through multiple post-transcriptional and post-translational processes, including intracellular folding, processing, trafficking and extracellular assembly, and therefore increased intracellular collagen does not necessarily translate into increased secretion over 48 hours^42,54,62^. Alternatively, these findings suggest that loss of PKN2 primarily promotes ECM remodelling rather than directly increasing collagen deposition. This interpretation is supported by increased secretion of MMP-1, enrichment of matrix remodelling pathways across both omics datasets, and impaired collagen fibre organisation observed in the zebrafish wound model.

Our findings also identify VEGF signalling as an important downstream pathway regulated by PKN2. Increased VEGF expression was observed at both transcript and protein levels following PKN2 depletion. Elevated VEGF has been implicated in fibroblast survival, angiogenesis and tissue remodelling, while VEGF signalling is therapeutically targeted by nintedanib, one of only three approved antifibrotic therapies for IPF^6,48,52^. Together with enrichment of ECM, adhesion and WNT pathways, these findings suggest that reduced PKN2 promotes tissue remodelling through multiple complementary mechanisms rather than a single downstream effector.

Importantly, these molecular changes translated into altered tissue repair. PKN2 depletion accelerated wound closure in primary human fibroblasts despite no evidence of increased proliferation, suggesting that altered ECM organisation, rather than enhanced proliferative capacity, underlies this phenotype. Increased secretion of MMP-1 and VEGF, together with altered expression of collagen, integrins and WNT signalling components, supports a model in which loss of PKN2 promotes dynamic ECM remodelling rather than simply increasing matrix accumulation. This concept was further supported *in vivo*, where pharmacological inhibition of PKN disrupted epithelial organisation, altered collagen fibre alignment and modified wound repair dynamics in zebrafish larvae. Together, these complementary human and zebrafish studies demonstrate a conserved role for PKN signalling in coordinating epithelial and mesenchymal responses during tissue repair.

Our findings also provide functional context for emerging evidence implicating PKN2 in fibrotic disease. Allen *et al.* identified genetic variation at the PKN2 locus associated with accelerated decline in FVC in patients with IPF, implicating PKN2 in disease progression despite the underlying biology remaining unknown. This genetic association provides an additional line of evidence supporting PKN2 as a therapeutic candidate, consistent with studies demonstrating that drug targets supported by human genetics have a substantially higher probability of clinical success^63^. Although we did not observe an association between sputum PKN2 expression and lung function, PKN2 expression in sputum strongly correlated with epithelial cell abundance, suggesting that sputum-derived PKN2 mRNA primarily reflects epithelial cell content rather than fibroblast biology or disease severity. In contrast, our mechanistic studies in primary human fibroblasts demonstrate that reduced PKN2 promotes fibroblast activation, extracellular matrix remodelling and dysregulated tissue repair, providing a potential biological explanation for the GWAS association. Furthermore, a recent study identified PKN2 as a critical regulator of cardiac fibrosis and myocardial remodelling following injury, suggesting that PKN2 may represent a conserved regulator of tissue remodelling across multiple organs^24,64–66^. To our knowledge, this is the first study to define the epigenetic regulation and functional role of PKN2 in human pulmonary fibrosis, integrating human tissue analyses with epigenetic, transcriptomic, proteomic and complementary *in vivo* validation.

Interestingly, this contrasts with findings in the heart, where PKN2 has been reported to promote fibrotic remodelling following injury, suggesting that its function may be highly tissue- and context-dependent^66^. This divergence highlights the potential challenges of therapeutically targeting PKN2 systemically, as strategies aimed at restoring PKN2 expression in the lung could potentially have adverse effects on cardiac remodelling. Therefore, although PKN2 may represent a potential target for pulmonary fibrosis, further investigation of tissue-specific functions and approaches to selective pulmonary modulation will be required before its therapeutic potential can be established.

Several limitations should be acknowledged. Our mechanistic studies focused primarily on fibroblasts despite PKN2 being expressed across multiple pulmonary cell populations, and therefore additional cell-specific functions remain to be explored. Furthermore, although our findings implicate DNA methylation in regulating PKN2 expression, additional mechanisms, including chromatin accessibility, histone modifications and post-transcriptional regulation, are also likely to contribute. Finally, the zebrafish studies employed pharmacological inhibition of PKN signalling rather than PKN2-specific genetic manipulation, although the *in vivo* findings provide independent support for a conserved role of this pathway during tissue repair.

## Conclusion

In summary, we identify PKN2 as a previously unrecognised regulator of human lung fibroblast activation, ECM remodelling and tissue repair. Reduced PKN2 expression in IPF is associated with altered DNA methylation and is further suppressed by profibrotic TGF-β1-dependent signalling. Loss of PKN2 induces coordinated transcriptional, proteomic and functional reprogramming involving extracellular matrix, adhesion, WNT and VEGF pathways, resulting in altered collagen homeostasis and dysregulated matrix remodelling. In vivo, inhibition of PKN signalling disrupts tissue repair and collagen fibre organisation, providing evidence that PKN2 contributes to the regulation of fibroblast-mediated matrix architecture during wound healing. Together, these findings establish PKN2 as an important regulator of pulmonary fibroblast homeostasis and tissue repair. The opposing effects of PKN2 reported across tissues, including its pro-fibrotic role in cardiac remodelling, highlight the context-dependent nature of PKN2 signalling and the need for further investigation of its tissue-specific functions before therapeutic modulation can be considered.

## Acknowledgements

The authors acknowledge the use of the NIHR Leicester Biomedical Research Centre (BRC) Immunohistochemistry (IHC) facility and the University of Leicester Integrated Histopathology and Imaging Core (IHIC) facility, including use of the Zeiss microscopy facilities. The authors thank Dr Kees Straatman for their technical assistance and support.

## Author Contributions

K.M.R., R.J.A., and L.V.W. contributed to the conception and design of the project. C.E.M. performed most of the experiments, analysed and interpreted the data, and contributed to manuscript preparation. O.A.P. designed and performed the fish experiments and contributed to data analysis, interpretation and manuscript preparation. P.B., V.M., B.G., and L.C. contributed to the collection of clinical sputum samples. P.R., A.V., P.S., G.E., M.W., M.R., S.N., A.J.H., H.M., B.L., D.J.L.J., R.L.C., and C.B.M. contributed to experimental work, data generation, analysis, and/or interpretation. K.M.R. oversaw and supervised the project, contributed to data analysis and interpretation, and wrote the manuscript. All authors discussed the results and commented on the manuscript.

