## Supplementary Materials for "Loss of PKN2 drives fibroblast reprogramming and extracellular matrix remodelling in pulmonary fibrosis"

**Supplementary Materials and Methods:**

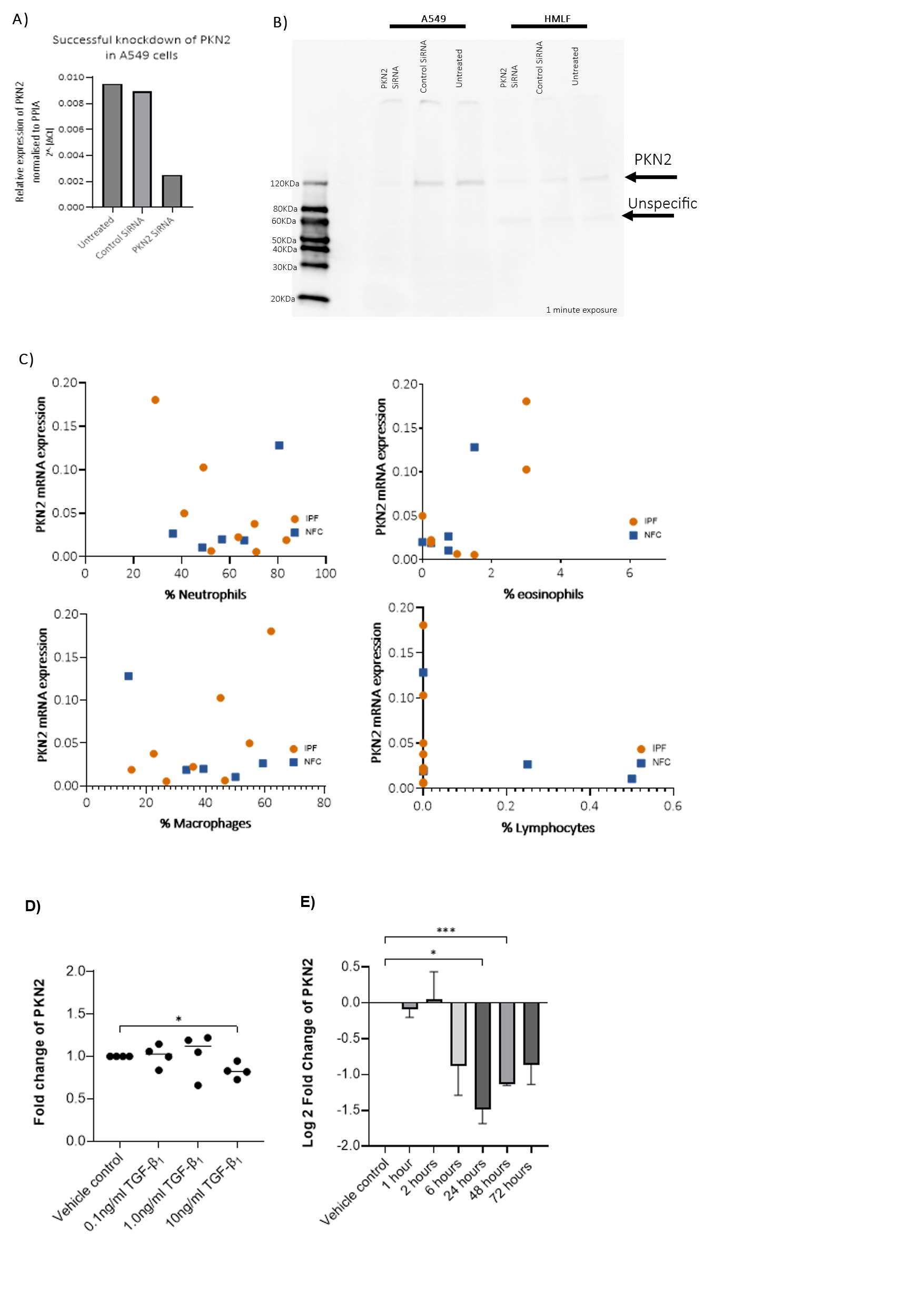

**Supplementary Figure 1. Validation of PKN2 expression analyses and supporting experimental data.**

**A)** qRT-PCR confirmation of efficient PKN2 knockdown following siRNA transfection in A549 epithelial cells (n=1). PKN2 expression was normalised to PPIA.

**B)** Western blot validation of the anti-PKN2 antibody (Thermo Fisher Scientific, MA5-15887). siRNA-mediated knockdown confirmed that the immunoreactive band at approximately 112 kDa corresponds to PKN2 in both A549 cells and HLMFs, whereas a lower molecular weight band remained unchanged following knockdown, indicating non-specific antibody binding in fibroblasts.

**C)** Correlation analyses demonstrating no significant relationship between sputum PKN2 mRNA expression and the proportions of neutrophils, eosinophils, macrophages or lymphocytes.

**D)** Dose-response analysis showing the effect of TGF-β1 (0.1, 1 and 10 ng/ml) on PKN2 expression in NFC fibroblasts. Maximum suppression of PKN2 expression was observed following treatment with 10 ng/ml TGF-β1 (n = 3).

**E)** Time-course analysis of PKN2 expression following stimulation with 10 ng/ml TGF-β1, demonstrating sustained downregulation of PKN2 over 72 hours (n = 3). Repeated measures ANOVA. *P < 0.05, **P < 0.01, ***P < 0.001.

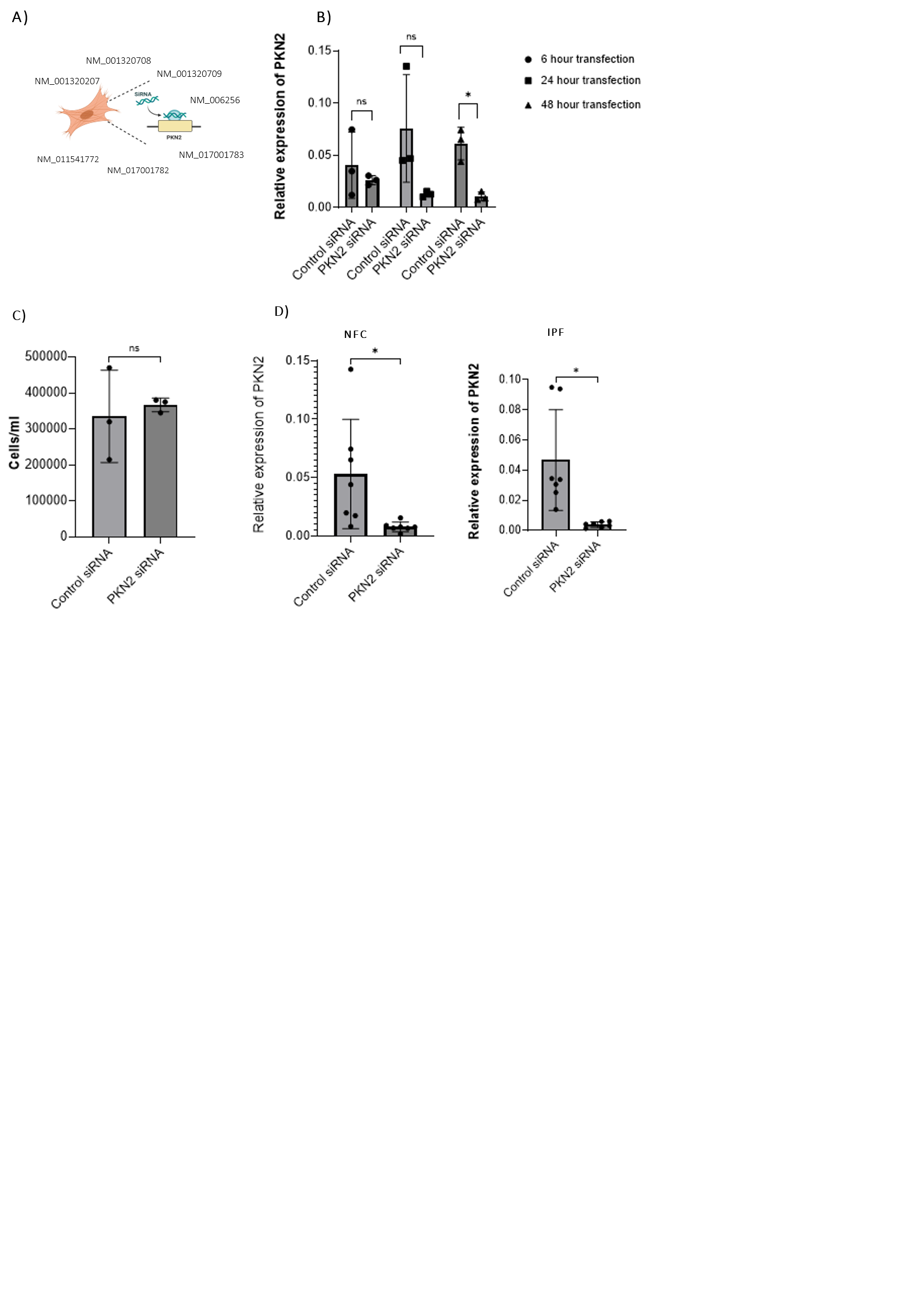

**Supplementary Figure 2. Validation and optimisation of siRNA-mediated PKN2 knockdown in primary human lung fibroblasts.**

A) Schematic illustrating the human PKN2 transcript variants targeted by the SMARTpool siRNA (Dharmacon).

B) Optimisation of siRNA-mediated PKN2 knockdown in non-fibrotic control (NFC) fibroblasts. qRT-PCR analysis demonstrates efficient suppression of PKN2 expression following 6, 24 and 48 h transfection with pooled PKN2 siRNA (50 nM) relative to non-targeting control siRNA (n = 3). Expression was normalised to PPIA.

C) Cell viability analysis showing that 48 h transfection with PKN2 siRNA does not alter NFC fibroblast cell number compared with non-targeting control siRNA, as assessed by trypan blue exclusion (n = 3).

D,E) Validation of siRNA-mediated PKN2 knockdown in primary fibroblasts isolated from NFC (n = 7)(D) and IPF n = 7 (E) donors. qRT-PCR confirmed significant reduction in PKN2 mRNA expression following 48 h transfection with pooled PKN2 siRNA. Statistical analysis was performed using paired t-tests.

**Materials and Methods:**

**Human tissue collection**

Lung tissue from NFC and IPF donors was collected under the following ethics: 10/H0402/12, 22/EM/0159 and 07/MRE08/42. NFC samples were collected from patients who were undergoing lung resection surgery for carcinoma at Glenfield Hospital, tissue removed was non-fibrotic tissue. IPF samples were collected at Glenfield Hospital.

**Immunohistochemistry staining of lung tissue**

Paraffin embedded sections of NFC and ILD lung tissue were examined for PKN2 expression using a PKN2 antibody (ThermoFisher Scientific, #MA5-15877) at a dilution of 1:1000. A ready to use isotype control was also used. Images were analysed using QuPath (v0.2.0-m2) software [1] to determine the percentage of positive cell staining.

**Sputum**

Sputum collection

Sputum was induced in NFC and IPF donors. Sputum plugs were isolated and processed using PBS and 0.2% DTT. Cell pellets were collected, reconstituted in RNAprotect before storage at -80°C until RNA isolation was performed. Sputum samples were collected from NFC and IPF donors under the ethics 10/H0402/12.

Sputum processing

Sputum plugs were isolated and weighed before PBS was added to plugs (at a volume of 8x weight) and rocked on ice for 15 minutes. Sputum plug in PBS was centrifuged and supernatant removed. Then 0.2% DTT was added to remaining sample at a volume of 4x weight of sputum before vortexing and rocking on ice for 15 minutes. Sample was then passed through 48µm gauze in a funnel into a clean falcon tube before weighing. A total cell count was performed with 10µl sample and 10µl trypan blue using a haemocytometer. Viable, dead and squamous cells were counted. The remaining sample was centrifuged and supernatant removed, cell pellet was resuspended in PBS and centrifuged. The cytospin slide was prepared and centrifuged, before supernatant was removed and cell pellet reconstituted in RNAprotect. The RNA protect mixture was stored at -80°C until RNA isolation was performed.

**Fibroblasts**

Fibroblast collection

NFC fibroblasts were derived from healthy areas of lung from patients at Glenfield Hospital, Leicester, UK who were undergoing lung resection for carcinoma. No morphological evidence of disease was found in the tissue samples used for fibroblast isolation as described previously [2, 3]. Fibroblasts were characterised using as described previously [2].

IPF derived fibroblasts were obtained from patients who were undergoing lung biopsy for diagnostic purposes at the University of Pittsburgh Medical Centre, USA (Dr Carol Feghali Bostwick: Department of Medicine, Division of Pulmonary, Allergy, and Critical Care Medicine, University of Pittsburgh, Pittsburgh, Pennsylvania, United States of America), other IPF fibroblasts were isolated from post mortem lung tissue of IPF patients (Dr Amanda Tatler: University of Nottingham, Nottingham, United Kingdom) and from Royal Papworth Hospital (Cambridge, United Kingdom).

Fibroblast cell culture

Fibroblasts were cultured in 10% FBS DMEM which contains: Dulbecco’s Modified Eagles Medium (DMEM), high glucose with GlutaMAX™ supplement (Gibco, #61965059), 1% antibiotic/antimycotic mixture (AA) (Gibco, #15240062), 1% non-essential amino acids (NEAA) (Gibco, #11140035) and 10% fetal bovine serum (FBS) (Gibco, #A5256801). Fibroblasts were maintained in 10% FBS DMEM media and incubated at 37°C with 5% CO_2_. Fibroblasts in culture were passaged using 0.25% trypsin-EDTA (Fisher Scientific, #11570626) to detach and HBSS to wash. All fibroblasts used were at latest passage 7.

Prior to some TGF-β1 stimulation experiments, fibroblasts were washed twice with Hanks’ Balanced Salt Solution (HBSS) (Fisher Scientific, #11540476) and then placed into serum free DMEM for 24 h before experimental conditions were applied. Serum free DMEM consists of: DMEM high glucose with GlutaMAX™ supplement (Gibco, #61965059), 1% AA mixture (Gibco, #15240062) and 1% NEAA (Gibco, #11140035).

Mycoplasma testing

Isolated NFC and IPF fibroblasts were tested for mycoplasma using the MycoStrip® mycoplasma detection kit (InvivoGen, #rep-mys-50) following manufacturers protocol. If negative, fibroblasts could be immediately used for experiments. If fibroblasts tested positive for mycoplasma infection, treatment with 12.5 µg/ml Plasmocin® (InvivoGen, #ant-mpt-1) in 10% FBS DMEM was carried out for 14 days before re-testing. If fibroblasts remained positive following this initial treatment period they were disposed of, if testing negative after this 14-day period fibroblasts could be used for experimental purposes.

**RNA isolation**

Lung tissue

RNA was isolated from homogenised lung tissue using the RNeasy Fibrosis Tissue Mini Kit (Qiagen, #74704) by the automated QIAcube connect system (Qiagen) following manufacturer’s instructions.

Fibroblasts

RNA was isolated from NFC and IPF fibroblasts using the RNeasy Plus Mini Kit (Qiagen, #74134) and QIAshredder columns (Qiagen, #79654). RNA isolation was carried out following the manufacturer’s instructions supplied with the kit and 50µl of RNA was collected for each sample.

Sputum

Following collection of sputum and differential cell counting, sputum was reconstituted in RNA Protect and stored at -80°C until RNA isolation was performed. RNA was isolated from NFC and IPF sputum samples using the PicoPure™ RNA isolation kit (ThermoFisher Scientific, #KIT0204). RNA isolation was carried out following the manufacturer’s instructions supplied with the kit and 13 µl of RNA was eluted for each sample.

RNA concentration and purity was quantified using the NanoDrop™ 2000 spectrophotometer (Thermo Scientific).

**Stimulation with Transforming Growth Factor Beta-1 (TGF-β1)**

For dose response and time-course experiments, fibroblasts from NFC and IPF donors were cultured to 80% confluency before being placed in serum free DMEM for 24 h. Fibroblasts were stimulated with either vehicle control or TGF-β1 at concentrations of 0.1 ng/ml, 1.0 ng/ml and 10 ng/ml (Peprotech, #100-21) for the stated experimental time. Fibroblasts were then washed with HBSS, trypsinised and collected before RNA was isolated using the method described above.

**siRNA mediated knockdown of PKN2 in fibroblasts**

Isolated fibroblasts were transfected with 50 nM siRNA SMARTPool targeting PKN2 (Dharmacon™, #M-004612-03-0005), or 50nM of a non-targeting control siRNA (Dharmacon™, #D-001210-03-05). In a 6-well plate, isolated fibroblasts were plated at 200,000 cells per well in 10% FBS DMEM and incubated overnight. The following day the culture media was removed and replaced with transfection mixture. The transfection mixture consisted of serum free DMEM, INTERFERin® transfection reagent (Polyplus, #101000028) and either PKN2 siRNA or non-targeting control siRNA. This transfection mixture was added to the fibroblasts with 10% FBS DMEM and incubated for the period stated (6, 24 or 48 h). RNA and supernatants were collected at the end of the 48-hour experimental period.

**Cell viability**

To examine the effect of PKN2 siRNA on cell viability, the trypan blue exclusion method was used. At the end of the experiment, fibroblasts were detached using 0.25% trypsin-EDTA, neutralised with 10% FBS DMEM and centrifuged to obtain a cell pellet. This was then reconstituted in 10% FBS DMEM, then a cell count was performed using 10µl cell suspension and 10µl trypan blue via a haemocytometer.

**Methylation experiments**

DNA methylation profiles of cultured airway and lung parenchymal fibroblasts from healthy and IPF patients (Healthy – 8 airway and 14 lung parenchymal fibroblasts; IPF – 8 airway bronchial and 8 lung parenchymal fibroblasts) was profiled using the Illumina HumanMethylationEPIC BeadChip array. Briefly, genomic DNA was extracted from each sample using the AllPrep DNA/RNA Mini Kit (Qiagen) according to the manufacturer’s instructions. DNA concentration and purity were assessed with a NanoDrop™ 8000 Spectrophotometer (Thermo Fisher Scientific). A total of 750 ng of DNA per sample was subjected to bisulfite conversion using the EZ DNA Methylation Kit (Zymo Research), following the supplier’s protocol to allow the identification between methylated and unmethylated cytosines. 160 ng of converted DNA was processed for genome-wide DNA methylation profiling. The bisulphite-converted DNA is fragmented into smaller pieces and then hybridized to the array.

HumanMethylationEPIC BeadChips were scanned using an Illumina HiScan system, and raw IDAT files generated in GenomeStudio were imported into R (2020) with the minfi package (v1.52.1) (Aryee et al., 2014; Fortin et al., 2017). Initial quality control identified no samples requiring exclusion based on median signal intensity, sex discordance, or beta distribution outliers. Functional normalization was applied using preprocessFunnorm in minfi (Fortin et al., 2014). Probes were filtered to remove those with low bead count, SNP overlap, poor detection p-values (≥0.01), sex chromosome mapping, cross-hybridization, or non-CpG targets (Pidsley et al., 2016). 730,159 probes remained for downstream analysis. Batch effects related to chip ID and sample position were corrected sequentially using ComBat in the sva package (v3.54.0) (Leek et al., 2012. Both beta values (proportion of methylation over total intensity) and M values (ratio of methylated to unmethylated intensity) were derived for interpretation and statistical testing respectively. The Beta values of methylation sites annotated to PKN2were subset and differential methylation analysis was performed through linear regression modelling using limma package (v3.62.1).

**RNA extraction and Gene Expression Microarray**

Total RNA was isolated using the AllPrep DNA/RNA Mini Kit (Qiagen) as per the manufacturer’s instructions. The quality and quantity of RNA were assessed using the NanoDrop™ 8000 Spectrophotometer (Thermo Fisher Scientific). RNA was reverse transcribed to produce cDNA, which was used as a template to generate cRNA. The cRNA was amplified, fragmented and labelled with a fluorescent dye. The labelled cRNA was then hybridized to different probes on the Affymetrix Genechip Human Gene 2.1 ST arrayand washed to remove any non-specific binding. The chip was then scanned using a high-resolution scanner and the fluorescent signal corresponds to the amount of bound cRNA to each probe. The raw data was converted to numerical intensity values which represent gene expression values, and these cel files were used for analysis.

Raw CEL files were imported into R statistical software using Affy package (version 1.84.0) (Gautier et al, 2004) and subject to GC Robust Multiarray Average (GCRMA) normalisation using the gcrma package (version 2.78.0) (Gentry, 2020). Following quality control checks after normalisation, principal component analysis revealed two non-specific clusters that did not associate with any covariates which were adjusted for using the comBat function in SVA package (version 3.54.0) (Leek et al., 2012). Gene expression of PKN2 was subset from the pre-processed dataset and plotted in Prism Graphpad.

**DNMT Inhibitor (5aza) Experiment**

Human lung fibroblasts were cultured to approximately 75% confluence in 6 well plates in serum positive DMEM. At ~75% confluence media was changed for fresh serum positive media containing 0.0001 µM, 0.001 µM, 0.01 µM, 0.1 µM, and 1.0 µM, 10.0 µM or 20.0 µM of 5-aza (5-Aza-2′-deoxycytidine, Sigma-Aldrich, #189825). Media only and DMSO was used as the vehicle control. 48 h later cells had reached confluence, and the media was replaced with serum free media containing 5-aza at the above concentrations. Following 24 h serum starvation the media was changed a final time and cells were maintained in 5-aza for a further 24 h. Following treatment, cells were lysed directly in-well using RLT buffer (Qiagen) supplemented with 10% β-mercaptoethanol. Genomic DNA and total RNA were simultaneously isolated using the AllPrep DNA/RNA Mini Kit (Qiagen). Lysates were homogenized and applied to AllPrep spin columns to sequentially purify genomic DNA and RNA. DNA and RNA were eluted in buffer EB and RNase-free water respectively. Nucleic acid concentration and purity were assessed using NanoDrop™.

0.5 μg of RNA was reverse transcribed using SuperScript IV Reverse Transcriptase (ThermoFisher Scientific) as per the manufacturer’s instructions. The resulting 20  μl cDNA samples were diluted to a total volume of 200  μl using nuclease-free water. cDNA was amplified using PerfeCTa SYBR Green FastMix (Quanta bio). qPCR was then performed using gene specific primers to measure expression of PKN2 (Sigma-Aldrich). PKN2 forward (sequence CGGAACTGCAGGGGGATTC) and PKN2 reverse (TCTGCTGCACCATTGTATCTGA). qPCR was carried out using a AriaMX qPCR System (Agilent Technologies).

**Quantitative real-time polymerase chain reaction**

cDNA was synthesised from lung tissue, fibroblast and sputum RNA using SuperScript IV VILO Master Mix (ThermoFisher Scientific) or LunaScript® RT SuperMix Kit (New England Biolabs) according to manufacturing instructions using the ProFlex PCR System (Applied Biosystems, Life Technologies). qRT-PCR was performed using gene specific TaqMan primers to measure expression of PPIA and PKN2 (ThermoFisher Scientific). Primer details are included in Table 1. qRT-PCR was carried out using a QuantStudio 5 Flex Real-Time PCR System (ThermoFisher Scientific).

| **Gene** | **ID** |
| --- | --- |
| PPIA | Hs99999904_m1 |
| PKN2 | Hs00178944_m1 |

**Table 1:** *TaqMan™ Gene expression assays used for qRT-PCR analysis.*

**RNA sequencing**

RNA sequencing was performed on NFC fibroblasts (n = 5) transfected with either 50 nM PKN2 siRNA or 50nM non-targeting control siRNA . Samples underwent quality control checks, RNA library prep and were sequenced using Illumina PE150 technology. Sequencing was performed externally by Novogene (Novogene UK Company Limited, Cambridge, United Kingdom).

**Proteomics**

Cell pellets were collected from NFC fibroblasts (n = 3) following siRNA transfection and the proteome was examined using bottom-up proteomics by liquid chromatography-mass spectrometry (LC/MS) as described below.

In-solution digestion and clean-up using SP3

Cell pellets were lysed by sonication (5 min) in 500 μL RIPA buffer (Thermo Scientific, Loughborough, UK) with complete mini protease inhibitor cocktail (Roche, Welwyn, UK). Protein concentration was determined by BCA assay and 10 μg protein per sample was randomised across a twin.tec 96-well protein LoBind plate (Eppendorf, Hamburg, Germany). Samples were processed by automated single-pot, solid-phase-enhanced (SP3) sample preparation[4] using an Agilent Bravo 96LT liquid handling system[5]. Reduction and alkylation were performed in a single step using 10 mM TCEP and 40 mM chloroacetamide in 50 mM ammonium bicarbonate pH 7.4 at 95 °C for 5 min. Proteins were bound to a 50 μg/μL suspension of 1:1 hydrophilic and hydrophobic Sera-Mag SpeedBeads (Cytiva Life Sciences, Buckinghamshire, UK) in the presence of acetonitrile, followed by washes with 80% ethanol and acetonitrile. Proteins were digested with trypsin/LysC (Promega, Southampton, UK) at a 1:25 enzyme-to-protein ratio for 16 h at 37 °C with agitation (350 rpm). Peptides were acidified with TFA to 1% final concentration and eluted from the beads.

Shotgun liquid chromatography-mass spectrometry

Peptide concentrations were determined by o-phthalaldehyde (OPA) assay and 600 ng peptide per sample was loaded onto Evotips for analysis by LC-MS/MS using the Evosep ONE LC (Evosep Biosystems, Odense, Denmark) coupled to a timsTOF HT mass spectrometer (Bruker Daltonics, Bremen, Germany). Data processing was performed using ProteoScape version 2025b and Spectronaut version 20.0.250606.92449.

Proteomics LC-MS and Raw Data Processing Parameters

The LC was operated with a 100 SPD standard 11.5 minute analytical gradient using the EV1064 Endurance analytical column ReproSil-Pur C18, 3 µm beads by Dr Maisch, 8 cm x 100 µm. DIA-PASEF scan mode was used with the scan range set at 400-1000 *m/z*, the TIMS mobility range to 0.64-1.45 V cm^2^, and ramp and accumulation times were both set to 100ms. The method included 8 DIA-PASEF scans with three 24 Da windows per ramp, resulting in an estimated cycle time of 0.95 sec. Data was acquired using Compass HyStar version 6.3 with real-time processing in ProteoScape version 2025b. Raw data were processing using Spectronaut version 20.0.250606.92449 (Biognosys, Zurich, Switzerland)[6]. Database searching was performed against the *homo sapiens* reviewed FASTA using directDIA mode with BGS factory settings using enzyme cleavage rules for Trypsin/P, LysC.

**RNA sequencing and Proteomics Statistical Analysis**

All statistical analyses for RNA sequencing and proteomics were carried out using R version 4.5.0 (The R Foundation for Statistical Computing) on RStudio 2023.03.0 build 386 (RStudio, Inc., Boston, MA). Data pre-processing was carried out using the R packages NOISeq [7] and Caret [8] to filter features which were not statistically relevant from the dataset, normalise the data, impute zero values, and assess and remove sources of unwanted technical variation. Differential expression analysis was carried out using the R package Limma [9]. The results were filtered for proteins with Benjamini-Hochberg FDR-adjusted p-values <0.05 and log_2_(FC) > 0.58 (FC > 1.5). Metascape was used for gene ontology (GO) overrepresentation tests [10]. STRING (Search Tool for the Retrieval of Interacting Genes/Proteins) version 12.0 was used for functional protein association network analysis to predict protein-protein interactions [11]. Clustering of protein interactions was performed using the Markov Cluster (MCL) algorithm. Drug repurposing candidates were identified using the NeDRex plugin v2.1.0 in Cytoscape v3.10.2. A global network of disorder-disorder and gene-disorder association types were imported from NeDRexDB[12]. Following this, a custom set of seeds (the differentially expressed genes obtained from the multi-omics analyses) were uploaded. Using these specified seeds, drug repurposing candidates were identified and ranks derived by employing the TrustRank algorithm with the following parameters: include only direct and approved drugs, damping factor = 0.85, result size = 50.

**Enzyme linked immunosorbent assay**

Commercial sandwich enzyme linked immunosorbent assays (ELISAs) kits were used to measure the secretion of human pro-collagen 1α1 (R&D Systems, #DY6220-05), human VEGF (R&D Systems, #DY293B-05) and human total MMP-1 (R&D Systems, #DY901B-05) by the isolated fibroblasts into the culture supernatant. ELISAs were performed according to the manufacturing instructions.

**Scratch wound assay**

Isolated fibroblasts from NFC (n = 7) and IPF (n = 7) donors were plated into sterile 6-well plates and incubated with 10% FBS DMEM for 24 h before siRNA transfection as described above. After 24 h of transfection, three artificial wounds were induced into the cell monolayer using a 200 µl pipette tip, HBSS was used to wash and then fibroblasts were re-transfected with either PKN2 or non-targeting control siRNA for 48 h in 10% FBS DMEM. Images of the induced wounds were captured at 0, 24 and 48 h post-wound induction using the EVOS™ XL Core Imaging system with a 10x objective lens. Images were analysed on ImageJ. Three images were analysed per condition at each time point.

**Migration assay**

Isolated fibroblasts from NFC (n=3) donors were plated into sterile 6-well plates and incubated with 10% FBS DMEM for 24-hours before siRNA transfection as described above. After 24-hours of transfection, three artificial wounds were induced into the cell monolayer using a 200ul pipette tip, HBSS was used to wash and then fibroblasts were re-transfected with either PKN2 or control siRNA for 48-hours in 10% FBS DMEM. The plate was then incubated and imaged using the LiveCyte2 microscope system with images captured every 30 minutes. Three areas were imaged per condition at each time point. Images were analysed using the TrackMate Plugin on Fiji 1.54 to determine the max distance travelled vs Track mean speed before results were averages and analysed using GraphPad Prism.

**Zebrafish**

Zebrafish husbandry and lines used

Zebrafish work was conducted following procedures in accordance with the Animals Scientific Procedures Act 1986 (ASPA), the Animal Welfare Act 2006. All experiments were carried out with the ethical approval of the University of Bristol Animal Welfare and Ethical Review Body (AWERB).

Zebrafish (*Danio rerio*) stocks of transgenic and wildtype lines (Ekkwill) were maintained according to standard procedures in UK Home Office approved aquaria [13, 14]. Transgenic strains used were *Tg(krt19:tdTomato-CAAX)* [15] and *Tg(krt19:col1a2-GFP)* [16]. *Tg(krt19:col1a2-GFP)* transgenic adults used for collagen deposition experiments were maintained in the double mutant background *casper* [17], without melanocytes and iridophores, to allow the imaging of collagen in the wound. All zebrafish embryos and larvae were raised at 28.5 °C in 1x Danieau's in Petri dishes, and were staged according to Kimmel *et al*. [18]. Larval ages are expressed hours post-fertilization (hpf), and days post-fertilization (dpf).

Wounding assay and inhibitor exposure

Needle stick wounding was carried out as previously described [16, 19]. Briefly, 4 dpf larvae were anaesthetized in 2.5% v/v MS-222 (Sigma, #E10521), placed on a dish with a bed of 1% w/v agarose, and wounded under a stereoscope with a 30G hypodermic needle (BD Microlance #304000) on their left dorsal somite above the cloaca.

For pharmacological inhibition of PKN activity, immediately after wounding, 4dpf zebrafish larvae were incubated in 2 µM PKN1/2-IN-1 (MedChemExpress, #HY-145899), herein named PKNi, dissolved in 1x Danieau's, or in n% v/v DMSO, as control. Media with PKNi or DMSO was replaced daily.

Imaging

Before imaging, larvae were anesthetised with 0.003% MS-222 (Sigma, #E10521) and mounted in 1% w/v low-melting agarose (Sigma, #A9414) in 1x Danieau's soution in 35 mm glass-bottomed dishes (MatTek, #S319281) and covered with 1x Danieau's solution with 0.003% MS-222 anaesthetic. Confocal imaging was carried out using a Leica TCS SP8 AOBS confocal laser scanning microscope attached to a Leica DMi8 inverted epifluorescence microscope using a 40 x 1.3 NA objective. For collagen deposition experiments, a Leica SP8 AOBS confocal laser scanning microscope attached to a Leica DM6000 upright epifluorescence microscope with a 25 x 0.95 NA dipping objective was used.

Image analysis

Fiji version v1.54p [20], together with the plugin ModularImageAnalysis (MIA) [21], was used for image processing and analysis. We used custom pipelines created in MIA for image segmentation and analysis of melanocytes, keratinocytes, and collagen fibre alignment. During the morphometric analysis of keratinocytes, an ellipse was fitted to each cell after the segmentation of the apical surface of the keratinocytes. This was used to calculate the cell's apical aspect ratio, and for calculations of angle of cell elongation relative to the wound. In this case the longest axis of the fitted ellipse was used as elongation axis, and the angle to a straight line to the position of the wound was measured. The calculation of the alignment index of the collagen fibres in the wound was carried out as previously described [16].

**Zebrafish Statistical Analysis**

Statistical analyses were conducted in R [22]. For data visualisation the package ggplot2 was used [23]. Dotplots show mean ± 95 % confidence interval (CI) (package ggpubr [24]) and the statistical comparisons were carried out using the package rstatix [25].

**Statistical analysis**

GraphPad Prism (Version 9 and 10, GraphPad Software, San Diego, California, USA) was used for all data analysis (excluding RNA-sequencing and LC-MS proteomics). Data distribution and normality was assessed using the Shapiro-Wilk test. Exact statistical test used to analyse data was determined based on normality and selected test is stated in figure legend. Statistical significance was identified as P < 0.05. Statistical analysis methods used for RNA-Sequencing and proteomic analysis have been detailed above.

13.	Westerfield, M., *The Zebrafish Book. A Guide for the Laboratory Use of Zebrafish (Danio rerio)*. 5th ed. 2007.

22.	R Development Core Team, *R: A language and environment for statistical computing.* 2021, R Foundation for Statistical Computing: Vienna, Austria.

23.	Wickham, H., *ggplot2*. 2 ed. Use R! 2016: Springer Cham. XVI, 260.

24.	Kassambara, A., *ggpubr: 'ggplot2' Based Publication Ready Plots.* 2023.

25.	Kassambara, A., *rstatix: Pipe-Friendly Framework for Basic Statistical Tests.* 2023.
